# Ecological distribution, environmental roles and drivers of *Actinobacteriota* in two Mid-Atlantic estuaries

**DOI:** 10.1101/2025.11.21.689735

**Authors:** Mir Alvee Ahmed, Jojy John, Barbara J. Campbell

## Abstract

*Actinobacteriota*, a bacterial phylum renowned for members that produce bioactive compounds (e.g., antibiotics), has key roles in terrestrial and aquatic ecosystems. Although soil and marine/freshwater *Actinobacteriota* are well studied, functions and activities of their estuarine counterparts are poorly understood. We characterized 67 metagenome-assembled genomes (MAGs) belonging to 12 *Actinobacteriota* families from Chesapeake and Delaware Bay water samples across different seasons, salinities, and size fractions. MAGs from four dominant families, *Ilumatobacteraceae*, *Nanopelagicaceae*, *Microbacteriaceae*, and S36-B12, were examined in depth for their abundance, functional potential, estimated growth rates, and gene expression among samples. *Actinobacteriota* were most abundant in low- to medium-salinity samples during spring and summer. Their abundance patterns were strongly influenced by combinations of salinity, temperature, and phosphate, nitrate and silicate concentrations. Notably, many exhibited high estimated growth rates under low and medium salinities in summer. Members of the four major families showed a range of metabolic capacities from generalist to specialist, and all encoded biosynthetic gene clusters (BGCs) for secondary metabolites, particularly terpenes and betalactones, that were differentially expressed across conditions. Bay, salinity and size fraction were the primary drivers of gene expression differences. Distinct secondary metabolite genes were expressed between bays, with higher expression generally observed in medium compared to low salinities. These findings underscore the metabolic versatility and environmental responsiveness of *Actinobacteriota*, highlighting their active role in estuarine microbial communities and their contributions to biogeochemical cycling in dynamic coastal ecosystems.

**Importance:** Estuaries are dynamic transition zones where microbial communities play key roles in nutrient cycling and ecosystem functioning. This study provides new insights into the distribution of *Actinobacteriota* families and their responses to salinity and nutrient gradients in two major U.S. estuaries, the Chesapeake and Delaware Bays. We characterized the potential roles of *Actinobacteriota* in biogeochemical processes and secondary metabolite production, which may influence broad microbial interactions. These bacteria may also serve as a source of novel bioactive compounds, including antibiotics. By integrating ‘omic and environmental data, this work advances our understanding of *Actinobacteriota* adaptations and their ecology in estuarine environments.

## Introduction

Microorganisms produce diverse secondary metabolites serving complex functions, like communication and competition within microbial communities (1, 2). The phylum *Actinobacteriota* is well known for producing biologically and pharmaceutically relevant natural secondary metabolites both in soil and marine environments (3, 4). *Actinobacteriota* are abundant in soil, and freshwater ecosystems (5), and are also found in oceans, sediments, mangrove forests, marine animals, and plants as evidenced by cultivated isolates, 16S rRNA gene-based, and metagenomic analyses (6–10). Although typically considered as high-GC content soil bacteria, *Actinobacteriota* are also found in aquatic environments and are especially abundant in freshwater habitats (11–14). Small genome sizes and low GC content of freshwater *Actinobacteriota* led to the view that they are only distantly related to their terrestrial relatives with large genomes, and complex life cycles. However, Actinomycetes from both soil and freshwater provide many known clinical drugs (2, 15). While *Streptomyces* are known to produce two-thirds of all known natural antibiotics (16), the rate of new antibiotic discovery is low. Newer sources, like marine *Actinobacteriota,* are being explored for their ability to produce biologically active compounds including antitumor, antifungal, herbicidal and antiparasitic compounds. For example, marine *Salinispora* are a rich source of complex secondary metabolites that help define their ecological roles within specific niches, and many of these products possess drug-like potential (8). These findings highlight the promise of estuarine *Actinobacteriota* as novel bioactive metabolite sources.

Estuaries are unique transitional zones where freshwater and seawater mix, creating nutrient-rich environments that support high biological productivity (17, 18). The Chesapeake and Delaware Bays are geographically close, but differ in salinity gradients, freshwater input, turbidity and light penetration, nutrient availability, and physico-chemical parameters. These factors influence changes in microbial communities in these bays in different seasons and salinities (19–23). The moderately mixed Chesapeake and well-mixed Delaware Bay vary in the abundance of particles and attached microbial communities (24). Moreover, higher human impacts in Delaware Bay are expected to affect microbial composition and function (25).

Genome-resolved analyses of estuarine *Actinobacteriota* can reveal functions specific to environmental gradients and genes encoding secondary metabolites of potential impact on these microbial communities and their associated ecosystem. In this study, we analyzed *Actinobacteriota* metagenome-assembled genomes (MAGs) from two bays to assess their abundance, estimated growth rates, functional potential and activity through gene expression. Comparing predicted and actively expressed biosynthetic gene clusters (BGCs) across several draft genomes revealed how *Actinobacteriota* support diverse ecological strategies within different seasonal, spatial, and geochemical gradients.

## Results

### *Actinobacteriota* MAGs from the Delaware and Chesapeake Bays

Metagenomes and metatranscriptomes from the Delaware and Chesapeake Bay samples collected in different seasons, salinities, greater (G08, >0.8 μm) and smaller (L08, <0.8 μm) filter size fractions were sequenced independently, approximately representing particle-attached and free-living bacterial communities, respectively. The G08 size fraction may also include large or aggregated bacteria that are not attached to particles. Sample names like CPSum15L08 or DESum22DL08 have abbreviated bay names (CP or DE for the Chesapeake or Delaware Bay), followed by season (Spr, Sum or Fall for spring, summer or fall), numerical values for salinity in practical salinity units, PSU (06, 15 or 22 etc.), and size fractions (L08 or G08). When needed, D or N indicated day or night samples, respectively. MAGs have additional bin numbers after the metagenome name (e.g., CPSum15G08_bin_28).

In this study, 67 *Actinobacteriota* MAGs dereplicated at the species level (average nucleotide identity ≥95%) with >75% completeness and <5% contamination were analyzed. The average completeness and contamination of these MAGs were 85.64% ± 6.43 and 1.3% ± 1.08, respectively (Table S1). Their completeness ranged from 75.0% to 97.4%, with 28% of MAGs exceeding 90% and contamination remained low across most MAGs, ranging from 0% to 4.74%, with 82% of MAGs with contamination below 2%. *Actinobacteriota* estuarine MAGs had an average estimated genome size and GC content of 1.66 ± 0.55 Mb and 54.83 ± 9.62, respectively.

### Phylogenomics of *Actinobacteriota* MAGs

*Actinobacteriota* MAGs from this study belonged to three taxonomic classes: *Acidimicrobiia* (n=25), *Actinomycetia* (n=41), and *Thermoleophilia* (n=1) (Table S1). They belonged to 12 families where *Ilumatobacteraceae* (n=19), *Microbacteriaceae* (n=8), *Nanopelagicaceae* (n=18), and S36-B12 (n=12) were dominant (Fig. S1, Table S1). Occasionally, shortened family names were added to the MAG names for clarity (like Ilu_CPSum15G08_28, where Ilu signifies *Ilumatobacteraceae*). Average estimated genome sizes (in Mbp) of S36-B12 (2.14 ± 0.3) and *Ilumatobacteraceae* (1.91 ± 0.3) were significantly larger than those of *Microbacteriaceae* (1.2 ± 0.12) and *Nanopelagicaceae* (1.19 ± 0.2) (Kruskal-Wallis, c^2^ = 42.1, p < 0.001).

### In situ abundances of Actinobacteriota

The relative abundance (based on RPKG i.e., reads per kilobase per genome equivalent values) of *Actinobacteriota* MAGs and their corresponding transcripts, varied across different environmental conditions. *Actinobacteriota* were generally more abundant in summer compared to spring or fall, in both bays, particularly in the Delaware Bay samples (Fig. 1–2). Genomospecies belonging to the *Nanopelagicaceae* and some *Ilumatobacteraceae* were abundant in low salinity summer samples from both bays. In contrast, other *Ilumatobacteraceae* and *Microbacteriaceae,* as well as *Actinomarinaceae* genomospecies were abundant in medium and high salinity summer samples. Some MAGs of S36-B12 and *Microbacteriaceae* were relatively abundant in medium and high salinities in spring and summer, while a few were prominent mostly in spring medium and high salinity samples.

**Fig. 1.**
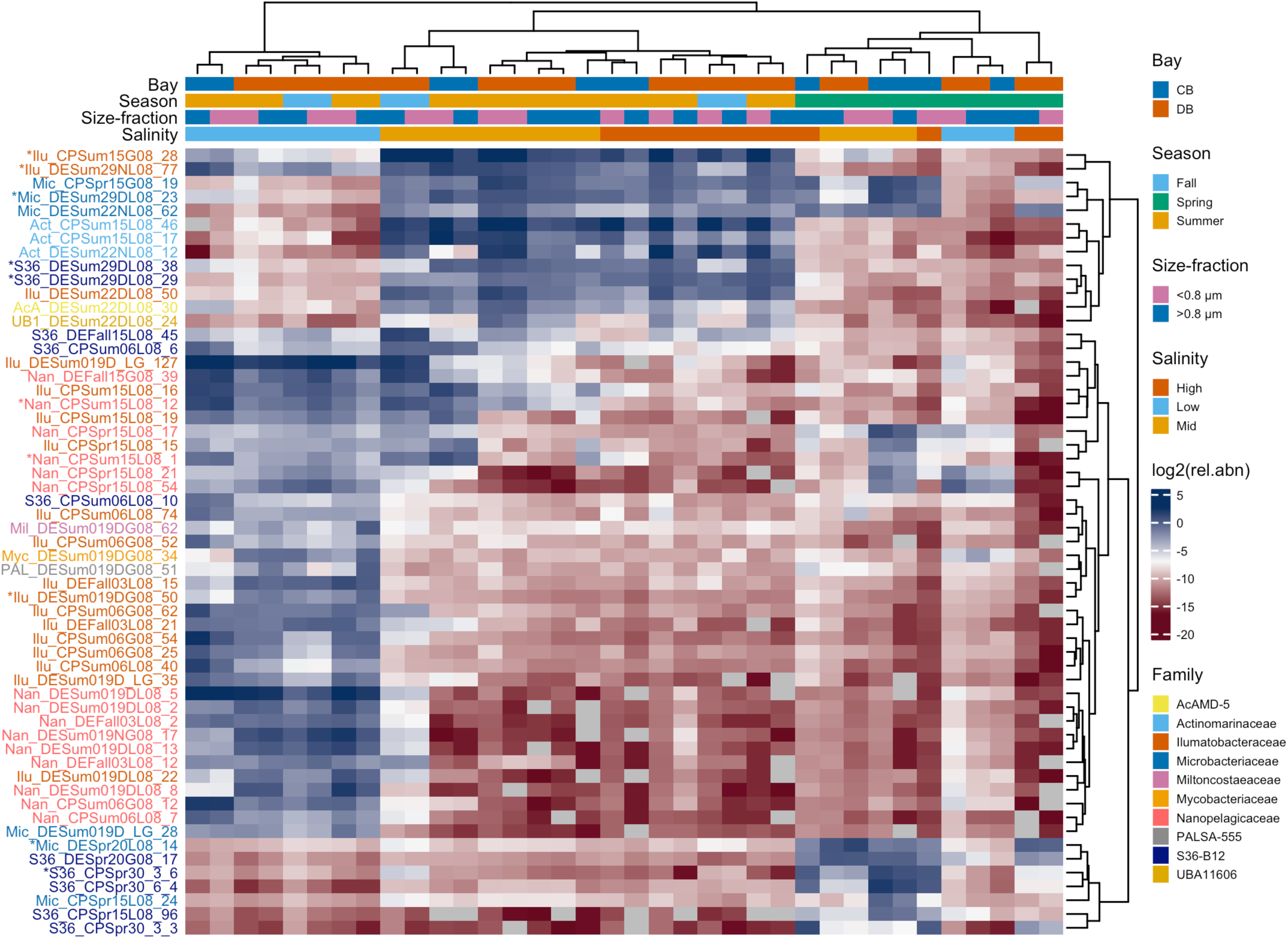
RPKG (reads per kilobase per genome equivalent) of *Actinobacteriota* genomospecies in the Chesapeake and Delaware Bay metagenomes. Boxes on the x-axis show bay, season, size-fraction and salinity of samples. Blue and red color cells of the heatmap indicate high and low relative abundance, respectively. Abbreviations: CP = Chesapeake Bay; DE = Delaware Bay; Spr = Spring; Sum = Summer; the numbers following indicate salinity in PSU; Hi = High; Lo = Low; and Mid = medium salinity; D = day; N = night; G08 = >0.8 µm, and L08 = <0.8 µm size fraction. Three letters at the beginning of a MAG indicate the genus to which it belongs according to GTDB-Tk taxonomy (100): AcA = AcAMD-5; Act = *Actinomarinaceae*; CAI = CAIXPF01; Ilu = *Ilumatobacteraceae*; Mic = *Microbacteriaceae*; Mil = *Miltoncostaeaceae*; Myc = *Mycobacteriaceae*; Nan = *Nanopelagicaceae*; PAL = PALSA-555; S36 = S36-B12; UB1 = UBA11606; UB5 = UBA5976. MAG names (*) and their families used in further analyses: Three MAGs (Ilu_CPSum15G08_28, Ilu_DEBay_Sum29NL08_77, Ilu_DEBay_Sum22DL08_50) from *Ilumatobacteraceae*, two (Mic_DEBay_Spr20L08_14, Mic_DESum29DL08_23) from *Microbacteriaceae*, two (Nan_CPSum15L08_12, Nan_CPSum15L08_1) from *Nanopelagicaceae* and three (S36_CPSpr30_3_bin_6, S36_DESum29DL08_bin_29, S36_DESum29DL08_bin_38) from S36-B12.

**Fig. 2.**
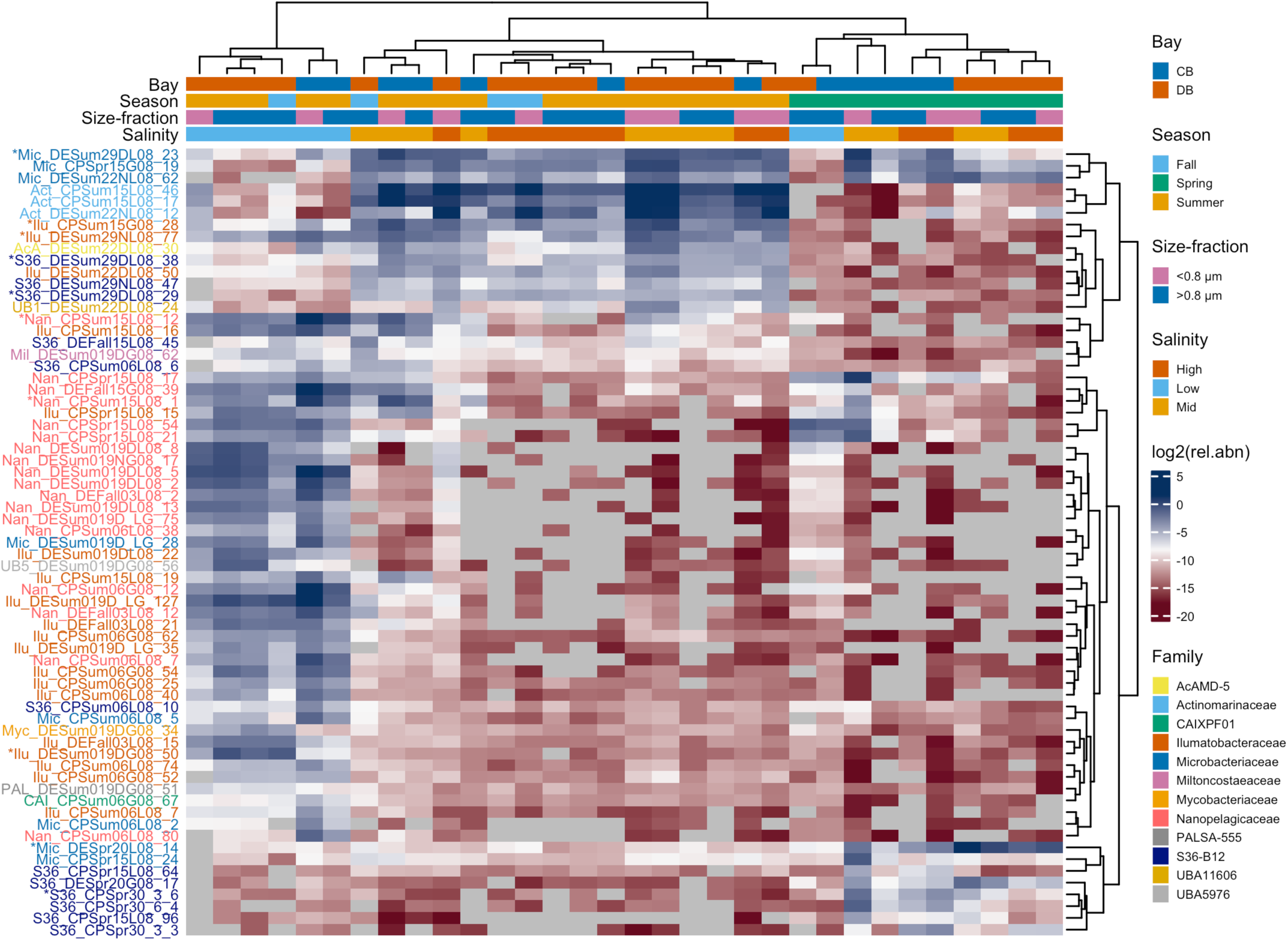
RPKG (reads per kilobase per genome equivalent) of *Actinobacteriota* genomospecies in the Chesapeake and Delaware Bay metatranscriptomes. Blue and red color cells of the heatmap indicate high and low relative abundance, respectively. Colors and abbreviations are same as Fig. 1.

Various combinations of environmental factors like salinity, nitrate and phosphate concentrations, temperature, chlorophyll *A* and silicate concentrations significantly affected the relative abundance of eight MAGs representing different *Actinobacteriota* family members in our samples (Fig. 3). Ilu_CPSum15G08_28 was significantly correlated with salinity and nitrate, while Ilu_DEBay_Sum22NL08_77 and Mic_DESum29DL08_23 were significantly correlated with nitrate and phosphate concentrations, and temperature. NanCPSum15L08_1 and Nan_CPSum15L08_12 were significantly correlated with salinity, chlorophyll *A*, and silicate concentration. S36_DESum29DL08_bin_29 and S36_DESum29DL08_bin_38 were significantly correlated with nitrate, phosphate concentrations, and temperature.

**Fig. 3.**
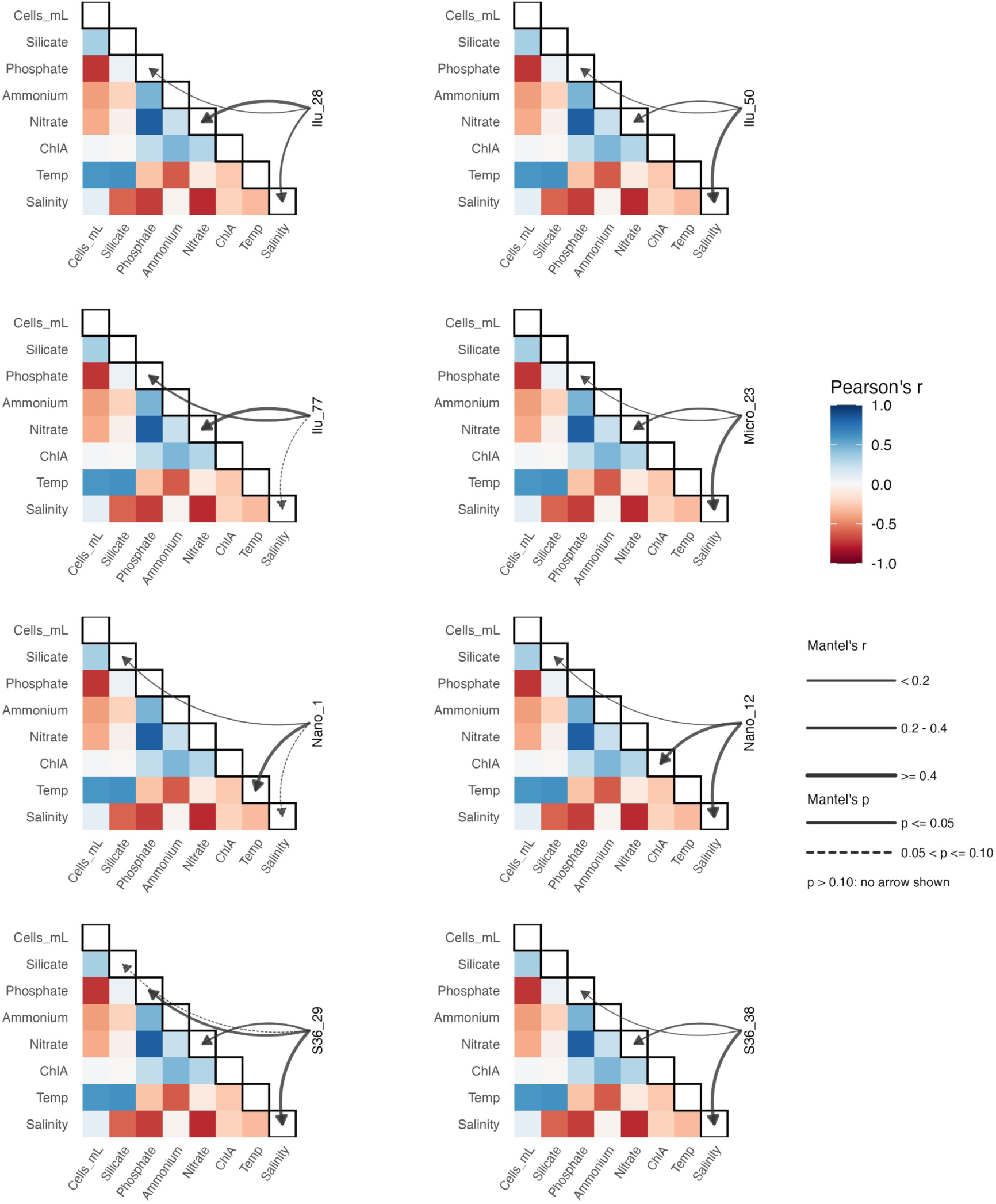
Mantel correlations between the relative abundance of representative Delaware and Chesapeake Bay *Actinobacteriota* MAGs and environmental factors. Abbreviations: Temp = Temperature; ChlA = chlorophyll *A* concentration; Cells_mL = cells per mL of water sample. Relative abundances (RPKG) of isolates: Ilu_28 = Ilu_CPSum15G08_28; Ilu_50 = Ilu_DEBay_Sum22DL08_50; Ilu_77 = Ilu_DEBay_Sum29NL08_77; Micro_23 = Mic_DESum29DL08_23; Nano_1 = Nan_CPSum15L08_1; Nano_12 = Nan_CPSum15L08_12; S36_29 = S36_DESum29DL08_bin_29; S36_38 = S36_DESum29DL08_bin_38.

### Metabolic potential of *Actinobacteriota*

*Actinobacteriota* MAGs were annotated to find general functional traits (e.g., central metabolism and transport) and specialized abilities (e.g., secondary metabolism and stress responses) in the genomes of the four major families. Comparisons of KEGG (Kyoto Encyclopedia of Genes and Genomes) (26) Orthology identifiers (KOs) between the four families revealed 616 shared metabolic and cellular functional traits, as well as distinct sets of unique functions in each (Fig. 4A, Dataset S1, Supplemental Text). For example, members of *Ilumatobacteraceae* and S36-B12 had more KO traits for energy and lipid metabolism than the other families (Fig. 4B). Moreover, compared to other three families, *Ilumatobacteraceae* exhibited more KO traits for xenobiotic compound degradation.

**Fig. 4.**
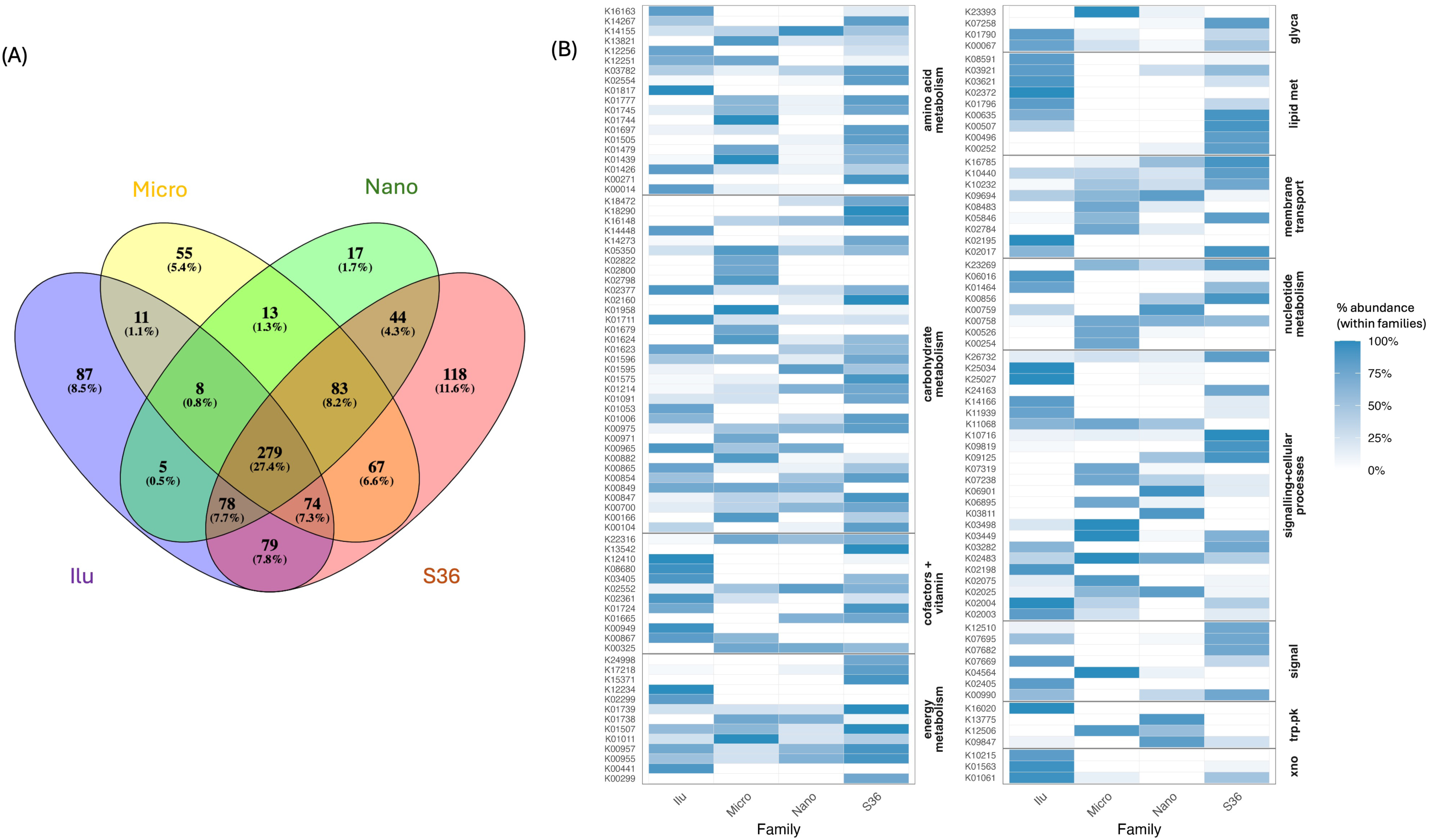
Functional traits across four *Actinobacteriota* families from the Delaware and Chesapeake Bays. Venn diagram showing the number of shared and unique KO traits across four families (A); Unique and shared KOs for MAGs within the four dominant families (B). Abbreviations: *lumatobacteraceae* (Ilu), *Microbacteriaceae* (Micro), *Nanopelagicaceae* (Nano), and S36-B12 (S36). Abbreviations: glyca = glycan metabolism; lipid met = lipid metabolism; signal = signal transduction; trp.pk = terpenoid and polyketide metabolism; xno = xenobiotic degradation.

*Ilumatobacteraceae* exhibited 87 unique KO traits, distributed across categories such as carbohydrate utilization, amino acid and fatty acid metabolism, terpenoid and polyketide biosynthesis (including ansamycin production), and xenobiotics degradation (fluorobenzoate, chloroalkane, and aminobenzoate) (Fig. 4, Dataset S1). Additional traits were associated with translation, protein folding and degradation, DNA replication and repair, and two-component signal transduction. Transport systems were also represented, including ABC transporters and quorum-sensing–associated transporters unique from other families.

*Microbacteriaceae* contained 55 unique KO traits spanning diverse categories including carbohydrate metabolism (glycolysis/gluconeogenesis, TCA cycle, and metabolism of fructose and mannose, galactose, ascorbate and aldarate, starch and sucrose, and propanoate), energy metabolism (oxidative phosphorylation, sulfur metabolism), nucleotide metabolism (purine and pyrimidine pathways), and amino acid metabolism (alanine, aspartate and glutamate, lysine, arginine and proline, and histidine) (Fig. 4, Dataset S1). Other unique traits were linked to glycan biosynthesis, cofactor and vitamin metabolism, terpenoid backbone biosynthesis, and transporters including ABC and LysE/ArgO.

*Nanopelagicaceae* displayed 17 unique KO traits related to carbohydrate, purine, amino acid metabolism, and terpenoid-associated functions (Fig. 4, Dataset S1). Distinct sugar and other (like *nodIJ*, *pnuC*) transport systems were also identified. Additional traits included a redox-sensitive regulator and DNA repair functions.

S36-B12 contained the most unique KO traits, 118, with broad representation across multiple categories. Carbohydrate metabolism included glycolysis, pentose metabolism, and starch processing, while energy metabolism was represented by sulfur pathways (Fig. 4, Dataset S1). Amino acid metabolism contained several biosynthetic components, and lipid metabolism was supported by additional traits. Transport systems were abundant, including both ABC-type and others. Distinct traits also included transcriptional regulators, ribosomal proteins, and stress-related functions.

### BGCs in Actinobacteriota

Varying numbers and types of BGCs in MAGs from the two bays were identified using three different annotation tools. First, various combinations of 15 BGCs were detected in 67 *Actinobacteriota* MAGs using antiSMASH (Fig. 5) (27). All MAGs belonging to *Microbacteriaceae* (n=8) and *Nanopelagicaceae* (n=18), and >75% (n= 9) and >61% (n=12) members of S36-B12 and *Ilumatobacteraceae*, respectively, had BGCs for terpene-encoding gene clusters. Betalactone-encoding gene clusters were found in three of the four major *Actinobacteriota* families except *Nanopelagicaceae*. Type III polyketide sulfate (T3PKS)-encoding genes were present in >33% members of S36-B12 only. Ranthipeptides-encoding genes were detected in >27% members of *Ilumatobacteraceae* only. Other BGCs were present in some members of the families, but only sporadically.

**Fig. 5.**
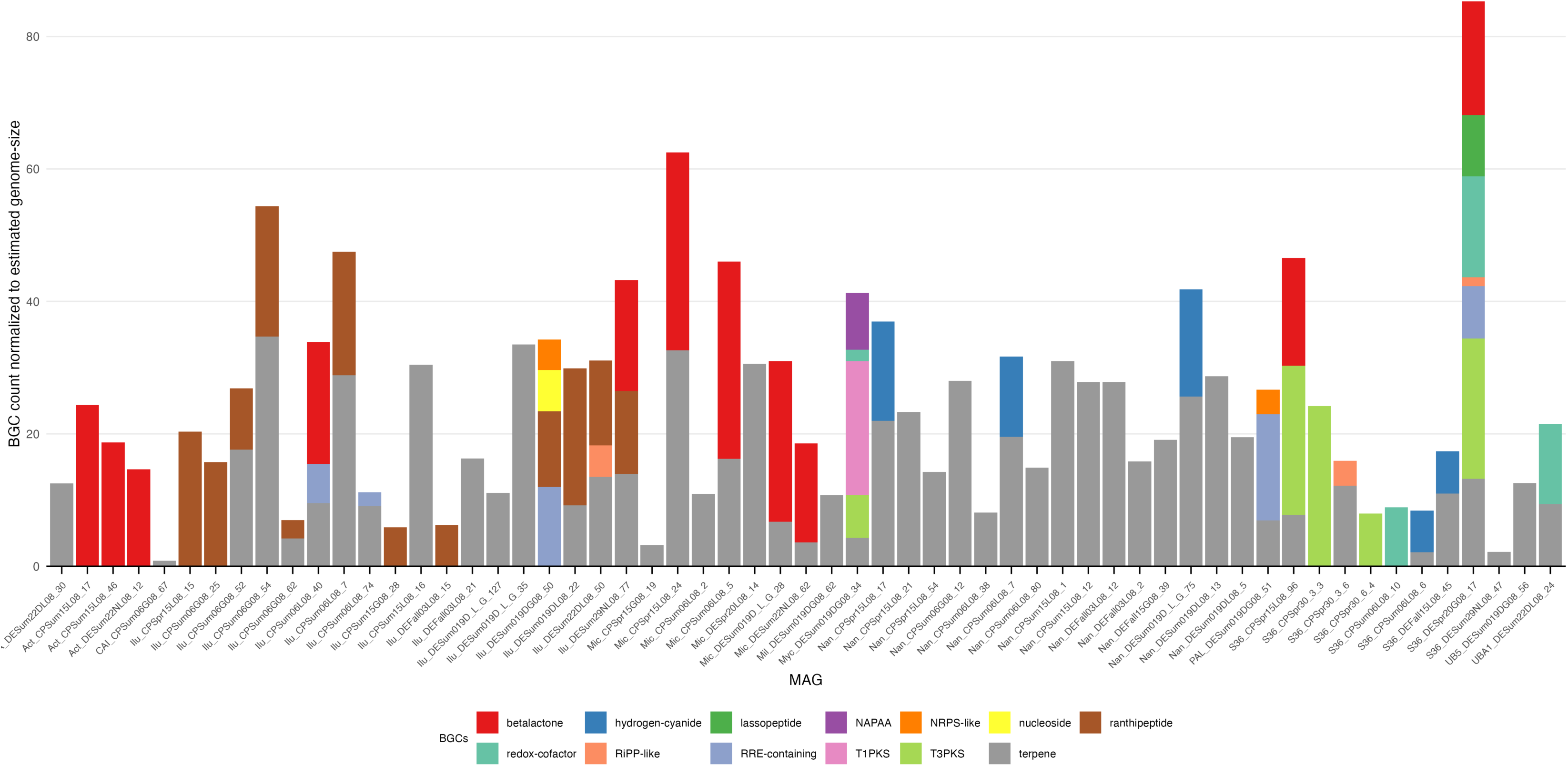
Biosynthetic gene clusters (BGCs) in *Actinobacteriota* MAGs from the Chesapeake and Delaware Bays. Numbers of BGCs detected in MAGs were normalized using a factor based on the smallest estimated genome size (∼0.77MBp). Abbreviations: RiPP-like = unspecified, ribosomally synthesized and post-translationally modified peptide products; T3PKS = Type III polyketide synthase; T1PKS = Type I polyketide synthase; NAPAA = non-alpha poly-amino acids (e.g., ε-polylysine); NRPS = non-ribosomal peptide synthetase; RRE-containing = cluster with RiPP Recognition Element. Three letters at the beginning of a MAG indicate the genus to which it belongs according to GTDB-Tk taxonomy (100): AcA = AcAMD-5; Act = *Actinomarinaceae*; CAI = CAIXPF01; Ilu = *Ilumatobacteraceae*; Mic = *Microbacteriaceae*; Mil = *Miltoncostaeaceae*; Myc = *Mycobacteriaceae*; Nan = *Nanopelagicaceae*; PAL = PALSA-555; S36 = S36-B12; UB1 = UBA11606; UB5 = UBA5976.

Second, using the keywords like “terpene”, “betalactones”, “resistance”, and “secondary metabolites”, multiple functions were listed from the InterProScan annotation output of 67 *Actinobacteriota* MAGs (28, 29). Compared to *Ilumatobacteraceae* or *Microbacteriaceae*, members of S36-B12, followed by *Nanopelagicaceae* showed more functions related to secondary metabolism (Fig. S2). Genes encoding glyoxalase for bleomycin/fosfomycin resistance, dienelactone hydrolase, and muconolactone delta-isomerase were uniquely detected in members of S36-B12. Genes encoding acriflavin resistance protein, and tellurite resistance methyltransferase were found in *Microbacteriaceae*. Moreover, secondary metabolite biosynthesis hydrolases, and copper resistance protein CopC/internalin were only seen in *Ilumatobacteraceae* and *Nanopelagicaceae*, respectively.

Third, Antibiotic Resistant Target Seeker (ARTS) was used for all MAGs and specially reported here for the ten for which RNASeq analyses were done. Ilu_DEBay_Sum22DL08_50 and Ilu_DEBay_Sum29NL08_77 each showed two BGCs for terpenes and betalactones, respectively (Dataset S1). Additionally, Nan_CPSum15L08_1, Nan_CPSum15L08_12, Mic_DEBay_Spr20L08_14, and Mic_DESum29DL08_23 each had one BGC for terpenes. S36-CPSpr30_3_bin_6 and S36-DESum29DL08_bin_29 contained two and one BGCs for bacteriocin, respectively. Additionally, S36-CPSpr30_3_bin_6, S36_DESum29DL08_bin_29 and S36_DESum29DL08_bin_38 all had one BGC for terpene.

### Estimated growth rates of *Actinobacteriota*

Analysis of peak-to-trough ratios (PTRs) (30) revealed contrasting patterns of estimated growth rates (log_2_PTR). Some *Ilumatobacteraceae*, *Microbacteriaceae*, *Nanopelagicaceae* and S36-B12 MAGs had variable log_2_PTR values in the medium to high salinity summer samples (Fig. 6A). For instance, three *Ilumatobacteraceae* MAGs had log_2_PTR values between 0.5–4.7. The average log2PTR values of Ilu_CPSum15G08_28 and Ilu_DESum29NL08_77 were comparable between the two bays, ranging from 0.51–0.6. In contrast, Ilu_DEBay_Sum22DL08_50 exhibited higher growth rates in the Delaware Bay (1.05 ± 0.06) than in the Chesapeake Bay (0.23 ± 0.04). Mic_DESum22NL08_62 showed higher log_2_PTR values in spring (0.4–0.7) compared to summer (0.2–0.5) whereas Mic_DEBay_Spr20L08_14 had high log_2_PTR values (0.6–1.2) in spring only. S36-B12 MAGs S36_CPSpr30_3_6 and S36_DESpr20G08_17 had estimated growth rates between 1.2 and 1.5, in medium and high salinity spring samples from Chesapeake Bay. However, S36_DESum29DL08_29 and had estimated growth rates between 0.8 and 1.1 medium and high salinity summer samples in the Delaware Bay. Finally, *Nanopelagicaceae* MAGs exhibited log₂PTR values (0.3–2.6) in low to mid-salinity samples.

**Fig. 6.**
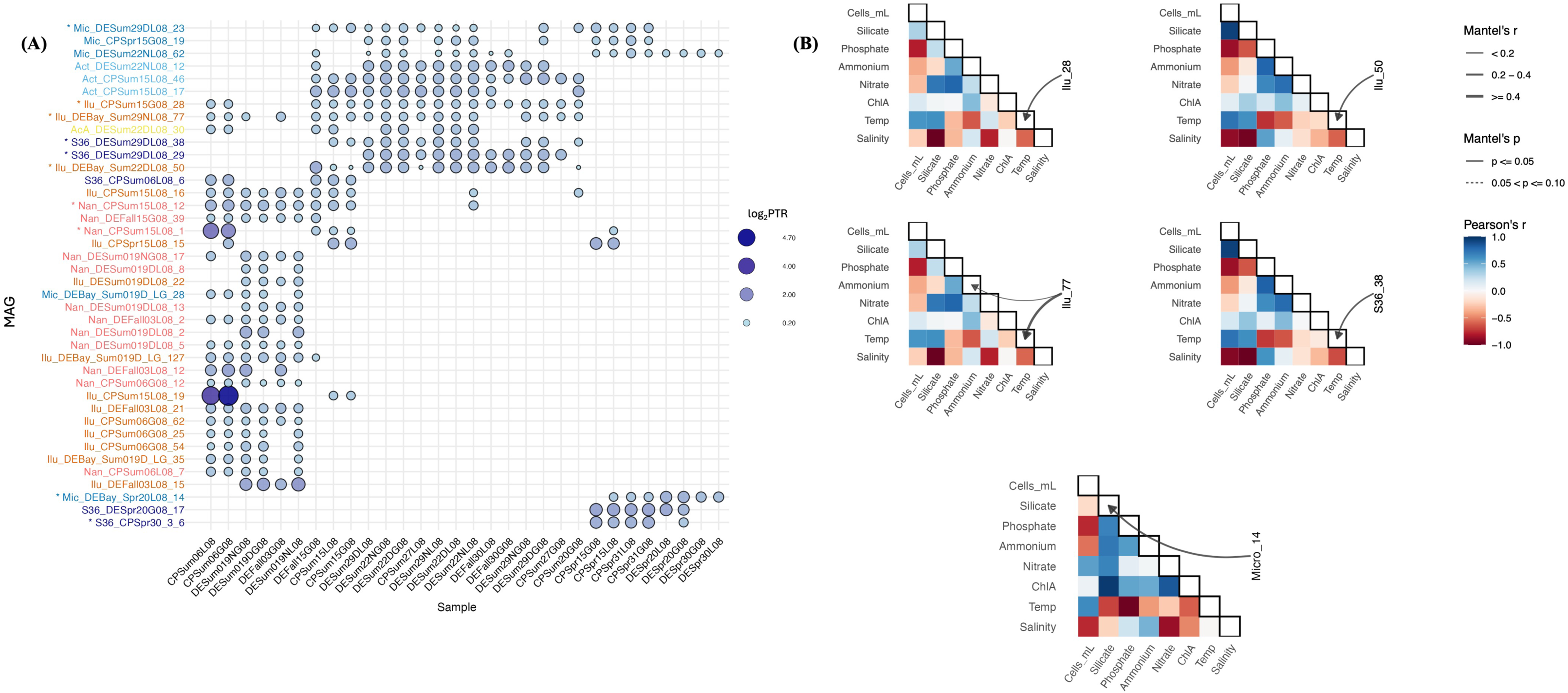
Estimated growth rates and environmental influence for *Actinobacteriota* in the Bays. **(A)** Estimated growth rates (log2PTR) of *Actinobacteriota* in the Chesapeake and Delaware Bays. **(B)** Correlation of PTR and environmental factors. MAG names with * were analyzed for differential gene expressions. Abbreviations: CP = Chesapeake Bay; DE = Delaware Bay; Spr = Spring; Sum = Summer; D = Day; N = Night; G08 = >0.8 µm; L08 = <0.8 µm size fraction. Abbreviations: AcA = AcAMD-5; Act = *Actinomarinaceae*; CAI = CAIXPF01; Ilu = *Ilumatobacteraceae*; Mic = *Microbacteriaceae*; Mil = *Miltoncostaeaceae*; Myc = *Mycobacteriaceae*; Nan = *Nanopelagicaceae*; PAL = PALSA-555; S36 = S36-B12; UB1 = UBA11606; UB5 = UBA5976. Ilu_28 = Ilu_CPSum15G08_28; Ilu_50 = Ilu_DEBay_Sum22DL08_50; Ilu_77 = Ilu_DEBay_Sum29NL08_77; Micro_14 = Mic_DEBay_Spr20L08_14; S36_38 = S36_DESum29DL08_bin_38.

Among the environmental factors examined, temperature significantly influenced estimated growth rates of four (Ilu_CPSum15G08_28, Ilu_DEBay_Sum22DL08_50, Ilu_DESum29NL08_77 and S36-DESum29DL08_bin_38) *Actinobacteriota* MAGs (out of the ten examined) (Fig. 6B). Estimated growth rates of Ilu_DESum29NL08_77 were also influenced by ammonium concentrations. In spring, estimated growth rates of Mic_DEBay_Spr20L08_14 were significantly correlated with silicate concentrations. However, none of the MAGs exhibited significant correlation with salinity, phosphate, nitrate or chlorophyl *A* concentrations.

### Changes in *Actinobacteriota* gene expression

Differentially expressed genes were observed from the ten MAGs mentioned above across samples under various salinities, seasons, locations (bay) and size fractions (Fig. S3). These factors strongly influenced differential gene expression in one or more of the above-mentioned MAGs (Table S2).

#### Differential gene expression between the two bays

Gene expression in *Actinobacteriota* MAGs varied between the two bays when controlling for other conditions, like in summer mid-salinity small size-fraction samples (Fig. 7, Dataset S2). *Ilumatobacteraceae* MAG Ilu_CPSum15G08_28 exhibited higher expression of several genes involved in carbohydrate metabolism in Delaware Bay than Chesapeake Bay. Increased expression of genes linked to branched-chain amino acid biosynthesis was also observed in the Delaware compared to Chesapeake Bay. Similarly, *Microbacteriaceae* MAG Mic_DESum29DL08_23 showed higher expression of multiple ABC-type sugar transporters, and glucitol/sorbitol PTS system genes and indirect CAZymes in the Delaware compared to Chesapeake Bay. This pattern was echoed by Mic_CPSpr15G08_14, which had more expression of sugar uptake genes in Delaware than Chesapeake Bay.

**Fig. 7.**
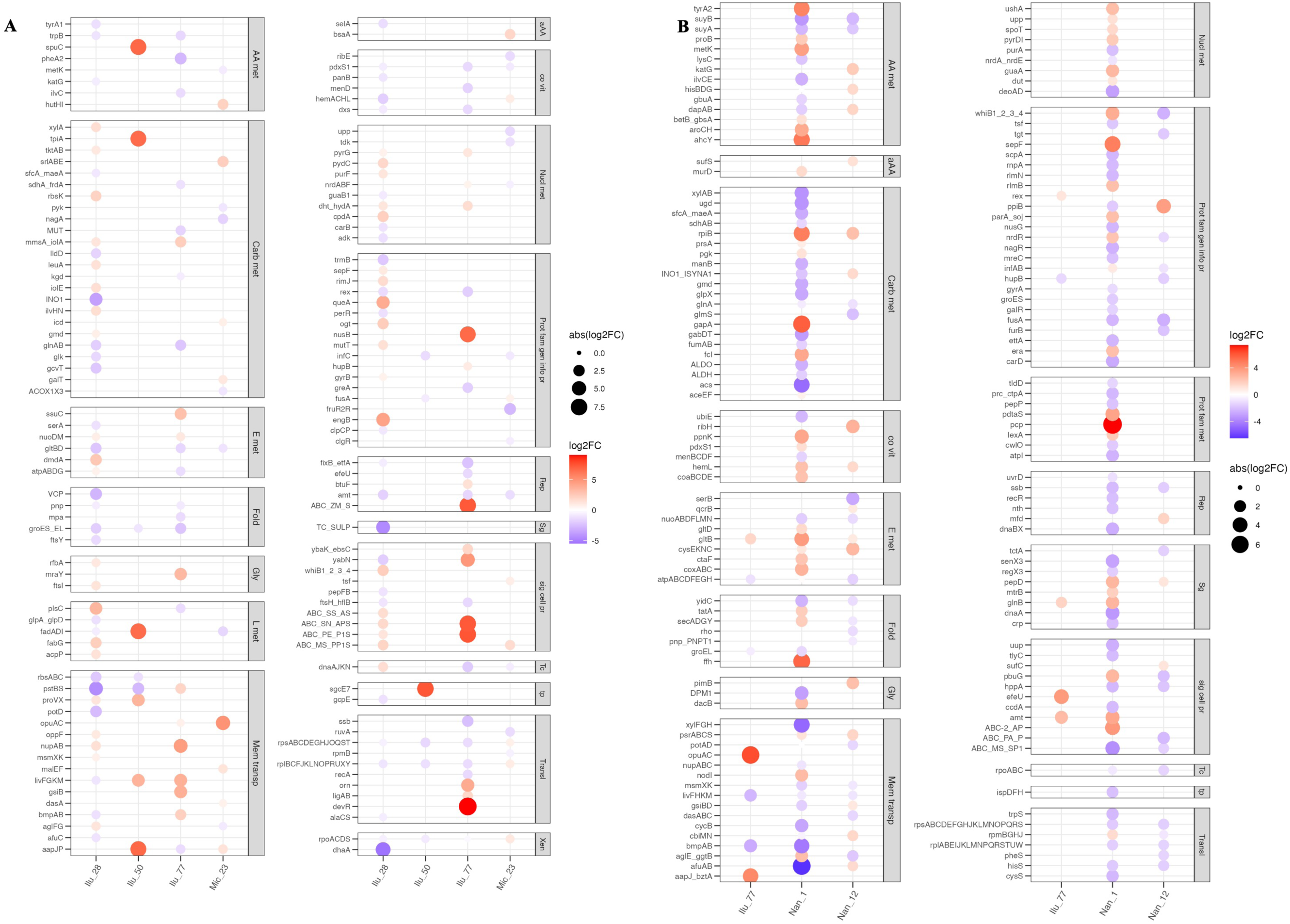
DESeq2 log2FoldChange values of significantly differentially expressed genes between **(A)** Delaware vs. Chesapeake Bays, and **(B)** Low vs. medium salinity environments. Red indicates higher expression in Delaware Bay or medium salinity. Absolute values of log2FoldChange are indicated by bubble size. Gene symbol abbreviations are mentioned in Dataset S2. Abbreviations: Ilu_28 = Ilu_CPSum15G08_28; Ilu_50 = Ilu_DEBay_Sum22DL08_50; Ilu_77 = Ilu_DEBay_Sum29NL08_77; Mic_23 = Mic_DESum29DL08_23; Nan_1 = Nan_CPSum15L08_1; Nan_12 = Nan_CPSum15L08_12.

Multiple ribosomal protein encoding and translation-related genes were more expressed in the Delaware than Chesapeake Bay across several genomospecies including Mic_DESum29DL08_23 and Mic_CPSpr15G08_14 (Fig. 7, Dataset S2). In contrast, Ilu_DEBay_Sum22DL08_50, Ilu_DESum29NL08_77 and S36_DESum29DL08_bin_38 had lower expression of genes for other ribosomal proteins and nitrogen regulatory proteins in the Delaware than Chesapeake Bay.

Other differentially expressed genes were involved in transport and oxidative stress response (Fig. 7, Dataset S2). Ilu_DESum29NL08_77 and S36-DESum29DL08_bin_38 showed higher expression of genes for glutathione transport, cobalamin, and osmoprotectants in the Delaware than the Chesapeake Bay. Expression of stress response genes such as glutathione peroxidase and thioredoxin-dependent peroxiredoxin were higher in Mic_DESum29DL08_23 in the Delaware compared to Chesapeake Bay.

Finally, several secondary metabolite biosynthesis pathways were differentially expressed between the bays (Fig. S4, Dataset S2). Ilu_DESum29NL08_77 exhibited higher expression of genes from betalactone and terpene biosynthetic clusters in Delaware than Chesapeake Bay, including three hypothetical proteins and one putative terpene-related gene. Conversely, in the same genomospecies, genes encoding a metal reductase, a D-3-phosphoglycerate dehydrogenase, and a disulfide formation protein showed significantly lower expression in Delaware than Chesapeake Bay. Moreover, S36_DESum29DL08_bin_29 expressed a gene for ubiquinone and other terpenoid-quinone biosynthesis significantly higher in the Delaware than the Chesapeake Bay. On the other hand, ribosomal proteins were expressed highly in the Chesapeake compared to Delaware Bay.

#### Differential gene expression between salinities

When we compared medium to low salinity small size fraction samples during summer from the Chesapeake Bay, five genomospecies showed significant differential expression of general and secondary metabolism genes (Fig. 7, Dataset S2). Several genes encoding ABC transporters showed contrasting expression patterns in Ilu_DESum29NL08_77. Genes involved in nitrogen metabolism, ammonium transport, and redox regulation were significantly more expressed in medium than low salinity. The genes encoding a DNA-binding protein and chaperone were expressed less under the same conditions.

We observed extensive differential gene expression between low and medium salinity conditions in two *Nanopelagicaceae* MAGs. In Nan_CPSum15L08_12, genes involved in oxidative phosphorylation, including multiple subunits of ATP synthase and cytochrome c oxidase showed significantly higher expression in medium than low salinity samples (Fig. 7, Dataset S2). In Nan_CPSum15L08_1, many genes related to transcription and translation showed significantly lower expression in medium compared to low salinity, including those encoding RNA polymerase subunits, multiple small and large ribosomal proteins, tRNA synthetases, and translation factors. Additionally, reduced expression was observed in transporter genes including sugar transporters, ion transporters, and peptide or amino acid transporters. In contrast, genes with higher expression in medium salinity than low salinity included those involved in amino acid biosynthesis, sulfur metabolism, fatty acid degradation, nucleotide metabolism, and porphyrin and vitamin biosynthesis. Several genes with functions in protein turnover and stress response, also showed higher expression under medium than low salinities.

Differential gene expression was also observed between medium and high salinities in two S36-B12 MAGs. In S36-CPSpr30_3_bin_6, ABC transporters involved in quorum sensing and phosphate transport, and genes for carbon monoxide dehydrogenation were expressed highly in medium than high salinity spring samples from the Chesapeake Bay. Additionally, genes for osmoregulation were expressed more in high salinity than medium salinity. In contrast, S36-DESum29DL08_bin_38 showed higher expression of genes for osmoprotection, ammonium transport and nitrogen regulation in high than medium salinity in the Chesapeake Bay summer samples.

In terms of secondary metabolism, distinct patterns emerged across three *Actinobacteriota* MAGs when comparing gene expression between medium and low salinities (Dataset S2). In Ilu_DESum29NL08_77, genes in betalactone and ranthipeptide biosynthesis clusters exhibited variable expression. Although most were not significantly different, one putative terpene biosynthesis gene had significantly lower expression in medium vs. low salinity, and another gene involved in disulfide formation also showed lower expression in medium vs. low salinity. In Nan_CPSum15L08_1, several terpene biosynthesis genes showed significantly more expression in medium vs. low salinities, including those involved in carotenoid production. On the other hand, Nan_CPSum15L08_12 showed an overall trend of lower expression of terpene biosynthesis genes in medium vs. low salinity samples, although none of the genes passed the significance threshold.

## Discussion

Our study provides a comprehensive view of the metabolism and activity of *Actinobacteriota* from the Chesapeake and Delaware Bays, highlighting their ecological roles and responses to estuarine environmental gradients. We found that temperature, salinity and phosphate, nitrate or silicate concentrations differently affected the abundance and estimated growth rates of representative MAGs within *Actinobacteriota* families. Moreover, potential metabolisms and realized differential expression of genes encoding general and secondary metabolites add to the existing knowledge about estuarine *Actinobacteriota*. Our findings offer new insights into the biogeography, functional potential, and activity of these bacterial populations, particularly concerning secondary metabolites.

### Ilumatobacteraceae

Consistent with a previous study of their dominance in the freshwater of the Delaware Bay (31), we found that *Actinobacteriota* were abundant in low salinity environments in both estuaries. Notably, MAGs belonging to *Ilumatobacteraceae* were consistently abundant in low to medium salinity summer samples of both bays, suggesting specialization to freshwater-influenced and transitional estuarine environments as described in other ecosystems (32–34). *Ilumatobacteraceae* also represent the most heterogenous *Actinobacteriota* family in Arctic fjords due to their abundance, genome makeup and functions (34).

Analyses of relative abundance and estimated growth rates of three *Ilumatobacteraceae* MAGs indicated that temperature influenced their active growth, but salinity shaped their distribution. This agrees with previously described 16S rRNA/rRNA gene estimates of abundance and presumed growth of some SAR11 species (31, 35). *Ilumatobacteraceae* were distributed and actively replicated in a range of environments, including medium (Ilu_CPSum15G08_28 and Ilu_DESum29NL08_77) and high (Ilu_DEBay_Sum22DL08_50) salinities, indicating their ubiquity. However, our correlation analysis was limited to low and medium salinity categories, which may have constrained detection of other environmental correlations. Functionally, this family exhibited capacities for oxidative phosphorylation, phosphonate transport, cytochrome P450 activity, and biosynthesis of vitamins and cofactors, indicating adaptation to redox-variable and nutrient-limited conditions.

*Ilumatobacteraceae* MAGs also encoded BGCs for terpenes, betalactones, and ranthipeptides, all with uncharacterized ecological functions in this lineage. However, studies in other marine bacteria have shown that these metabolites can provide competitive and defensive advantages, suggesting that these pathways represent promising targets for deeper ecological and biotechnological investigation (36–41). Terpenes are diverse in structure and bioactivity, often discussed as antimicrobial and cytotoxic agents (42, 43). Here, secondary metabolite encoding genes were differentially expressed across conditions, with some terpene transcripts more abundant in low than medium salinities, suggesting a role in microbial competition within a complex, terrestrially influenced freshwater community. Betalactone associated agents are involved in antimicrobial activities in natural environment and some are also used for clinical treatment (44). Widespread ranthipeptides, on the other hand, belong to the ribosomally synthesized and post-translationally modified peptide (RiPP) superfamily involved in quorum sensing (45). Genes for such compounds were commonly detected in *Ilumatobacteraceae* in agricultural soils (46). The presence of these genes suggests the potential of these bacteria to produce signaling or defense molecules to outcompete other community members.

Transcriptomic analyses further revealed environment-specific gene expression patterns among *Ilumatobacteraceae*. For instance, genes involved in nitrogen metabolism were expressed more in medium than low salinities, potentially reflecting the ecological distribution of Ilu_CPSum15G08_28 and Ilu_DESum29NL08_77, both of which showed strong correlations with nitrate and phosphate concentrations in addition to salinity. These patterns may support sustained physiological activity under transitional salinity conditions (47). Notably in Ilu_DESum29NL08_77, higher expression of genes related to stress response in medium than low salinity, and higher protein folding and ribosomal gene expression in low vs. medium salinity indicate a salinity-driven transcriptional shift. Also, Ilu_DEBay_Sum22DL08_50 exhibited significantly higher growth rates in the Delaware Bay than in the Chesapeake Bay together with lower expression of stress-related chaperones in the Delaware Bay. This suggests their elevated replication may co-occur with reduced stress response, possibly due to more favorable environmental conditions in the Delaware Bay.

### Nanopelagicaceae

*Nanopelagicacea*e were abundant in low salinity summer samples, and their abundance significantly correlated with silicate and chlorophyll *A* concentrations, suggesting a preference for freshwater and potentially phytoplankton-influenced conditions, as reported previously (48). This family is globally dominant in freshwater environments, often affected by phytoplankton abundance where they exhibit positive selection on genes involved in nucleic acid and amino acid metabolism in response to changing environmental conditions (48–50). As with other studies, members of this family showed streamlined genomes (51, 52). Our genome-resolved analyses revealed a broad functional repertoire including genes for sugar and amino acid metabolism, degradation of aromatic compounds, and quorum sensing, consistent with a generalist as well as an environmentally responsive lifestyle. Despite free-living lifestyles and streamlined genomes, their ability to use various nutrient-rich organic compounds was reported previously (49, 53, 54). All *Nanopelagicaceae* members from our study harbored terpene BGCs, particularly those associated with carotenoid biosynthesis which was suggested before as a fitness provider under light and dark, and as a protective agent against UV radiation (55, 56). Widespread distribution under low to medium salinity conditions highlighted their adaptive capacity to estuarine salinity gradients.

*Nanopelagicaceae* MAGs, Nan_CPSum15L08_1 and Nan_CPSum15L08_12, exhibited higher estimated growth rates in low than medium salinity samples; however, a statistically significant relationship between salinity and growth could not be confirmed, likely due to the limited number of samples. However, both MAGs had strong salinity-driven shifts in gene expression. For instance, both highly expressed genes in the nitrogen regulatory system in low compared to medium salinity, highlighting a common reliance on efficient nitrogen assimilation, a key adaptation in nutrient-variable, estuarine waters (50). Nan_CPSum15L08_1 also showed elevated ribosomal protein expression under low-salinity conditions reflecting enhanced growth relative to medium-salinity environments as ribosomal protein abundance often correlates with cellular growth rate (57). Additionally, their higher expression of oxidative phosphorylation, amino acid biosynthesis, and ABC transporter genes under medium compared to low salinities, pointed to a shift toward internal resource management and stress tolerance.

Nan_CPSum15L08_12 also showed reduced expression of transcription and translation genes under medium salinity than low salinity, suggesting a shift toward reduced protein synthesis and slower growth as observed in our and other previous studies (57, 58). This was accompanied by decreased expression of transporter genes, pointing to reduced nutrient uptake activity. However, in medium salinities, the higher expression of genes involved in amino acid biosynthesis, sulfur metabolism, fatty acid degradation, nucleotide metabolism, and vitamin/cofactor biosynthesis points to enhanced metabolic versatility or internal resource management compared to low salinities (50). Thus, both *Nanopelagicaceae* MAGs appear capable of reorganizing their physiology to balance growth, metabolism, and stress management across varying salinity conditions, a trend that aligns with observations from related MAGs in the Pearl River Estuary that exhibit high salinity tolerance (51). These strategies suggest niche partitioning within *Nanopelagicaceae* highlighting the importance of transcriptional plasticity in maintaining resilience under dynamic estuarine conditions.

### Microbacteriaceae

*Microbacteriaceae* genomospecies were enriched in medium to high salinity spring and summer samples from both bays suggesting their estuarine and marine preference compared to freshwater in our study, though in other studies they were abundant throughout a broad salinity range (59, 60). They demonstrated a multifaceted ecological strategy in estuarine systems, marked by flexible substrate utilization and environmental responsiveness. These bacteria appeared to adopt an opportunistic lifestyle, taking advantage of shifting nutrient regimes and salinity conditions. Similar results were observed previously with the freshwater- and brackish-dominant *Rhodoluna* and *Aquiluna Microbacteriaceae* isolates, respectively (61–64). *Microbacteriaceae* genomospecies in our study encoded and differentially expressed a diverse repertoire of carbohydrate metabolism-related genes, some indirectly related to CAZymes, enabling utilization of mannitol, rhamnose, ascorbate, and starch, substrates often associated with terrestrial input or phytoplankton-derived material. Previous genome- and isolation-based studies found various CAZymes in *Microbacteriaceae* from Antarctic and mangrove environments (64, 65).

Although not dominant in overall abundance or transcript levels, *Microbacteriaceae* demonstrated moderate to high estimated growth rates in spring and summer, implying responsiveness to seasonal nutrient availability, such as those following phytoplankton blooms. Notably, *Microbacteriaceae* were abundant in human impacted, nutrient rich regions of the Sydney Harbor (66). Also, in another study on the Powder River Basin, the abundance of *Microbacteriaceae* was higher in planktonic compartments compared to biofilm communities (60). However, our findings were also supported by positive correlations between *Microbacteriaceae* abundance and nitrate and phosphate concentrations, as well as between their estimated growth rates and silicate levels. Notably, like patterns observed in estuarine *Rhodobacterales*, the gene expression profiles of *Microbacteriaceae* also diverged between estuaries: transporter systems, carbohydrate metabolism (including fructose and galactose pathways), and oxidative-stress-related genes were highly expressed in Delaware than in the Chesapeake Bay (58). Moreover, a study on Columbia River Coastal Margin found that expression of such genes often shifts with nutrient regimes and competition along salinity gradients, reflecting differences in nutrient sources and potential microbial competition across these estuarine systems (67). Such transcriptional tuning suggests *Microbacteriaceae* can quickly adjust to changing environmental pressures or microbial interactions and this was particularly reported previously for soil environments (68, 69). The presence of terpene and betalactone biosynthetic gene clusters further implies a role in microbial interaction networks, possibly through chemical signaling or competitive inhibition (70, 71). Together, these traits and activities underscore the ecological significance of *Microbacteriaceae* as adaptable heterotrophs contributing to carbon processing, nutrient recycling, and community structure modulation in estuarine microbial assemblages. This was also evidenced from other studies where members of this family showed adaptation to environments contaminated with heavy metals, pharmaceutical waste or in complex interactions with plant roots (72–74).

## S36-B12

Members of the S36-B12 lineage were versatile, present in spring and summer, especially in medium to high salinity conditions, and covaried with salinity, nitrate and phosphate concentrations. Their larger genome size and high GC content compared to other estuarine *Actinobacteriota* was accompanied with many unique traits indicating their complex interactions with environmental factors. Members of this family were also reported with high GC content and mutation derived antibiotic resistance in the Caspian Sea (75). S36-B12 genomospecies encoded a rich suite of proteolytic enzymes, secretion systems, and phosphate acquisition genes, suggesting a role in recycling organic matter and coping with phosphate limitation during spring phytoplankton blooms. S36-B12 members were also found involved in sulfur cycling in Salton Sea, California (76). In summer, the contrasting expression profiles of S36-B12 indicated lineage-specific adjustments, where S36-B12_CPSpr30_3_bin_6 maintained metabolic activity at moderate salinity while DESum29DL08_bin_38 enhanced osmotic, nitrogen, protein folding and homeostasis responses to tolerate high-salinity stress in the Chesapeake Bay. Moreover, S36_DESum29DL08_bin_29 expressed secondary metabolite genes more strongly in the Delaware Bay and growth-related genes in the Chesapeake Bay, indicating a shift from stress adaptation to active growth across environments as observed in other *Actinobacteriota* families, *Rhodobacterales* and others (57, 58). Thus, we observed them harboring various BGCs and expressing genes differentially across bays and salinities highlighting their flexible salinity- and environment-responsive strategies along the estuarine gradient. Also, moderate growth rates of this family members, lack of correlation with any specific environmental factor and transcriptional data indicated low but detectable expression across conditions, hinting at a slow or more persistent growth strategy.

## Conclusions

This study revealed both estuary-specific ecological strategies and metabolic versatility of four families within the *Actinobacteriota*. Characteristics of the Chesapeake and Delaware Bays, particularly salinity, changing nutrient concentrations and temperature, emerged as key drivers of their lineage-specific distribution and activities. Moreover, their estimated growth rates significantly changed temporally, indicating seasonal increases in replication. BGCs for secondary metabolites, including terpenes, betalactones, and ranthipeptides were widespread in this phylum and differentially expressed, indicating important roles in microbial competition and environmental adaptation. The presence of these BGCs also suggests the potential to produce bioactive compounds with pharmaceutical relevance. Future studies should investigate the regulation of these BGCs under varying environmental conditions to evaluate their functional potential and applicability in drug discovery. The presence of oxidative stress genes and putative antibiotic resistance traits further highlights the capacity of these bacteria to thrive under fluctuating estuarine conditions. Overall, our findings from investigating this group with metagenomes and metatranscriptomes underscore the ecological significance of *Actinobacteriota* in estuaries.

## Materials and Methods

### Sampling

Surface water samples from the Chesapeake and Delaware Bays were collected and processed as previously described (77, 78). This included sequential filtration to separate particle-attached and free-living microbial fractions, followed by metagenomic and metatranscriptomic sequencing and analysis.

### Quality assessment and taxonomic affiliation of *Actinobacteriota* MAGs

Size, contig numbers, GC content and N50 value of MAGs were recorded through Quast v5.0.2 (79). CheckM v1.1.3 (lineage_wf default parameters) was used to find completeness and contamination of MAGs (80). Representative, dereplicated *Actinobacteriota* MAGs were picked using dREP (v3.4.0) based on 95% ANI (81). Phylogenetic relatedness of genomes/MAGs were analyzed, and a tree was made with GTDB-Tk v2.1.1 (classify_wf default parameters) (82). Tree visualization and annotation was done with iTOL v7.2.2 (83).

### *In situ* relative abundance and growth rates

Each metagenome and metatranscriptome were mapped to 67 dereplicated MAGs using Bowtie2 v2.4.5 with default parameters. Resulting sam files were converted into bam and sorted bam files with SAMtools v1.15.1 (84, 85). FeatureCounts v2.0.1 summarized read mapping results and then normalized relative abundances of each of 67 dereplicated *Actinobacteriota* MAGs were calculated based on the RPKG approach in each Chesapeake and Delaware Bay sample as previously described (86). A heatmap was made with ComplexHeatmap v2.24.1 library of R/RStudio v4.5.1 (87–89). To evaluate relationships between MAG relative abundance patterns and environmental variables, we applied one-sided Mantel tests (n = 999 permutations) using Bray–Curtis dissimilarity for MAG abundance and Euclidean distance for environmental data. Analyses were conducted separately for Summer samples, and significant correlations (p ≤ 0.1) were visualized in multipaneled correlation–scatter plots generated in R v4.5.1 using ggplot2 and patchwork (88, 90).

The ratio of sequence coverage of the origin and terminus of replication were obtained using the program Compute PTR (CoPTR) (30). PTR values were a proxy to understand estimated replication of representative *Actinobacteriota* MAGs in various conditions (30, 91) A bubble plot was prepared from this data using the ggplot2 package in RStudio (88, 92). To find correlation between estimated growth rates and environmental variables, one-sided Mantel tests in ggplot2 in R v4.5.1 was used (88, 90).

### Functional potential of *Actinobacteriota*

Prokka was used to annotate member MAGs of *Actinobacteriota* families (93). Then amino acid sequences were exported to BlastKOALA from KEGG Web service to find KO number assignment to the query gene. These KO numbers were compared across *Actinobacteriota* families to find common and unique traits. (26) Additionally, using AntiSMASH and InterProScan, we examined the presence of BGCs, and genes associated with antibiotic resistance, respectively, to better understand their ecological roles through producing secondary metabolites in the estuary (27, 94). Moreover, ARTS was used to find number of core genes, BGCs and BGC proximity in selected MAGs (95)

### Expression of BGCs

Number of metatranscriptome reads mapped against representative *Actinobacteriota* MAGs with Bowtie2 v2.4.5 (84) were used in count tables from featureCounts of subread package v.2.0.1 and were condensed based on gene clusters identified in Prokka (96, 97). Gene ids were further matched with KEGG defined KO numbers, and antiSMASH and InterProScan outputs (26, 28, 29, 98). Metatranscriptomes with at least ∼10,000 reads mapped to genes within a representative MAG were used for analysis in DESeq2 (99). From these count tables, principal component analysis (PCA) plots were generated to find sample clusters regarding gene expression. For each representative MAG, differentially expressed genes were identified using subsets of feature count tables created based on temporal or spatial differences of samples. To determine differentially expressed genes, adjusted p-values were used from DESeq2 analyses (99). The most highly expressed genes from few MAGs were used for visualization with DESeq2 and pheatmap packages in R (88, 93, 99)

## Data availability

The metagenomes, metatranscriptomes, and MAGs for this study are available on NCBI under the umbrella project PRJNA432171.

## Acknowledgements

Grants from the National Science Foundation (OCE-1261359 and EF-2025541) and the DOE (CSP-1621) to B.J.C supported this work. We acknowledge Clemson University for the generous allotment of compute time on the Palmetto Cluster supported by the National Science Foundation under Grant Nos. MRI# 2024205, MRI# 1725573, and CRI# 2010270. Moreover, the ‘omics analyses were done, in part, with the Clemson University Genomics and Bioinformatics Facility’s software support, through two Institutional Development Awards (IDeA) from the National Institute of General Medical Sciences of the National Institutes of Health under grant numbers P20GM146584 and P20GM139769.

We thank Dr. Matt Cottrell, Dr. Shen Jean Lim, and Liying Yu for help with sampling and nutrient analyses, Dr. Maximiliano Ortiz, Suzanne Crull, and Dinuka Lakmali Jayasuriya Patabandige for bioinformatic assistance and the crew of the R/V Hugh R. Sharp for help with sample collection. We also thank Tijana Galvina del Rio at the Joint Genome Institute for project help.

B.J.C. obtained funding for the study. B.J.C. supervised the project and directed sampling efforts. M.A.A. and J.J. analyzed the data and wrote the paper with input from B.J.C.

## Supplemental figure legends

**Fig. S1** Phylogenomics of *Actinobacteriota* MAGs from the Delaware and Chesapeake Bays. Clades with multiple genomes were collapsed into grey wedges where appropriate. MAGs from this study are shown in bold, and the twelve families containing these MAGs are colored. Abbreviations: CP= Chesapeake Bay; DE = Delaware Bay; Spr = Spring; Sum = Summer; the numbers following bay name and season indicate salinity in PSU; D = day; N = night; G08 = >0.8 µm, and L08 = <0.08 µm size fraction.

**Fig. S2** Antibiotic and heavy metal resistance genes in *Actinobacteriota* MAGs from the Chesapeake and Delaware Bays. Numbers of resistance genes detected in MAGs were normalized using a factor based on the smallest estimated genome size (∼0.77MBp). Abbreviations: ab = antibiotic; Enz = Enzyme; mdr = multidrug resistance; prot = protein; reg = regulator; res = resistance; Transp = Transporter; Three letters at the beginning of a MAG indicate the genus to which it belongs according to GTDB-Tk taxonomy (94): **AcA** = AcAMD-5; **Act** = *Actinomarinaceae*; **CAI** = CAIXPF01; **Ilu** = *Ilumatobacteraceae*; **Mic** = *Microbacteriaceae*; **Mil** = *Miltoncostaeaceae*; **Myc** = *Mycobacteriaceae*; **Nan** = *Nanopelagicaceae*; **PAL** = PALSA-555; **S36** = S36-B12; **UB1** = UBA11606; **UB5** = UBA5976.

**Fig. S3** Ordination plots of gene transcripts from seven abundant *Actinobacteriota* MAGs. PCA plot of DESeq2 normalized and transformed transcript abundances of representative MAGs from different families in different samples, corresponding to different environmental conditions. Abbreviations: CP = Chesapeake Bay; DE = Delaware Bay; Spr = Spring; Sum = Summer; the numbers following indicate salinity in PSU; Hi = High; Lo = Low; and Mid = medium salinity; D = day; N = night; G08 = >0.8 µm, and L08 = <0.8 µm size fraction. Three letters at the beginning of a MAG indicate the genus to which it belongs according to GTDB-Tk taxonomy: **Ilu** = *Ilumatobacteraceae*; **Mic** = *Microbacteriaceae*; **Nan** = *Nanopelagicaceae*.

**Fig. S4** DESeq2 log2FoldChange values of significantly differentially expressed BGCs between **(A)** Delaware vs. Chesapeake Bays. Red indicates higher expression in the Delaware Bay than the Chesapeake Bay. Blue indicates higher expression in the Chesapeake Bay than the Delaware Bay. Absolute values of log2FoldChange are indicated by bubble size. Meaning of gene symbols are mentioned in Dataset S2. Abbreviations: Ilu_50 = Ilu_DEBay_Sum22DL08_50; Ilu_77 = Ilu_DEBay_Sum29NL08_77.

**Fig. S5** DESeq2 log2FoldChange values of significantly differentially expressed BGCs between small (L08) and large (G08) size-fractions. Red indicates higher expression in L08 than G08 fraction and blue indicates the opposite. Absolute values of log2FoldChange are indicated by bubble size. Meaning of gene symbols are mentioned in Dataset S2. Abbreviations: Ilu_77 = Ilu_DEBay_Sum29NL08_77; Mic_23 = Mic_DESum29DL08_23; Nan_1 = Nan_CPSum15L08_1; Nan_12 = Nan_CPSum15L08_12.

**Table S1:** *Actinobacteriota* MAG quality and their taxonomic annotation.

| MAG name | CheckM Info |  | Quast outcome |  |  |  |  |  | GTDB-TK Taxonomy |  |  | RNA | CoPTR |
| --- | --- | --- | --- | --- | --- | --- | --- | --- | --- | --- | --- | --- | --- |
|  | Completeness | Contamination | # contigs | Largest contig | Total length | Genome Size (Mb) | GC (%) | N50 | Class | Order | Family |  |  |
| AcA_DESum22DL08_30 | 86.83 | 0 | 72 | 130644 | 1304696 | 1.30 | 42.03 | 73052 | Actinomycetia | Nanopelagiales | AcAMD-5 |  |  |
| Act_CPSum15L08_17 | 77.78 | 1.28 | 32 | 237889 | 1021372 | 1.02 | 31.13 | 79037 | Acidimicrobiia | Actinomarinales | Actinomarinaceae |  | Yes |
| Act_DESum22NL08_12 | 77.78 | 1.28 | 36 | 149773 | 1010635 | 1.01 | 34.07 | 57828 | Acidimicrobiia | Actinomarinales | Actinomarinaceae |  | Yes |
| Act_CPSum15L08_46 | 76.92 | 1.28 | 35 | 146843 | 997912 | 1.00 | 32.63 | 49481 | Acidimicrobiia | Actinomarinales | Actinomarinaceae |  | Yes |
| CAI_CPSum06G08_67 | 77.94 | 2.56 | 565 | 15146 | 1840162 | 1.84 | 71.48 | 3541 | Acidimicrobiia | Acidimicrobiales | CAIXPF01 |  |  |
| Ilu_CPSum15G08_28 | 97.44 | 0.85 | 277 | 65114 | 2123254 | 2.12 | 66.38 | 15352 | Acidimicrobiia | Acidimicrobiales | Ilumatobacteraceae | Yes |  |
| Ilu_CPSpr15L08_15 | 96.15 | 1.28 | 134 | 93811 | 1684427 | 1.68 | 52.04 | 28185 | Acidimicrobiia | Acidimicrobiales | Ilumatobacteraceae |  | Yes |
| Ilu_CPSum06G08_52 | 95.73 | 1.79 | 185 | 52786 | 1851906 | 1.85 | 53.75 | 14713 | Acidimicrobiia | Acidimicrobiales | Ilumatobacteraceae |  | Yes |
| Ilu_DEBay_Sum29NL08_77 | 94.02 | 0.85 | 35 | 383664 | 2230799 | 2.23 | 62.37 | 131726 | Acidimicrobiia | Acidimicrobiales | Ilumatobacteraceae | Yes |  |
| Ilu_CPSum15L08_19 | 93.16 | 1.28 | 127 | 72532 | 1659655 | 1.66 | 50.51 | 26362 | Acidimicrobiia | Acidimicrobiales | Ilumatobacteraceae |  | Yes |
| Ilu_DEBay_Sum019D_LG_127 | 93.16 | 0.43 | 92 | 164808 | 1542406 | 1.54 | 50.25 | 22844 | Acidimicrobiia | Acidimicrobiales | Ilumatobacteraceae |  | Yes |
| Ilu_DEBay_Sum22DL08_50 | 93.16 | 1.28 | 70 | 138438 | 2300829 | 2.30 | 63.87 | 60393 | Acidimicrobiia | Acidimicrobiales | Ilumatobacteraceae | Yes | Yes |
| Ilu_CPSum15L08_16 | 92.31 | 1.28 | 85 | 133641 | 1686707 | 1.69 | 61.42 | 41660 | Acidimicrobiia | Acidimicrobiales | Ilumatobacteraceae |  | Yes |
| Ilu_CPSum06G08_25 | 90.51 | 1.38 | 194 | 77099 | 1979814 | 1.98 | 68.36 | 16824 | Acidimicrobiia | Acidimicrobiales | Ilumatobacteraceae |  |  |
| Ilu_CPSum06G08_54 | 89.55 | 1.24 | 195 | 84186 | 1658229 | 1.66 | 61.64 | 16933 | Acidimicrobiia | Acidimicrobiales | Ilumatobacteraceae |  |  |
| Ilu_CPBay_Sum0_6L08_7 | 87.08 | 4.44 | 130 | 98171 | 1670203 | 1.67 | 58.9 | 18400 | Acidimicrobiia | Acidimicrobiales | Ilumatobacteraceae |  |  |
| Ilu_DEFall03L08_15 | 86.75 | 3.88 | 367 | 33585 | 1994675 | 1.99 | 65.25 | 6999 | Acidimicrobiia | Acidimicrobiales | Ilumatobacteraceae |  | Yes |
| Ilu_CPSum06G08_62 | 85.96 | 2.74 | 323 | 61355 | 2228781 | 2.23 | 62.62 | 10823 | Acidimicrobiia | Acidimicrobiales | Ilumatobacteraceae |  | Yes |
| Ilu_DESum019DG08_50 | 85.17 | 1.23 | 269 | 48099 | 2726823 | 2.73 | 65.71 | 13974 | Acidimicrobiia | Acidimicrobiales | Ilumatobacteraceae |  |  |
| Ilu_DEBay_Sum019D_LG_35 | 82.85 | 1.28 | 144 | 53409 | 1579293 | 1.58 | 60.03 | 15553 | Acidimicrobiia | Acidimicrobiales | Ilumatobacteraceae |  |  |
| Ilu_CPSum06L08_40 | 82.15 | 0 | 255 | 101347 | 2114545 | 2.11 | 65.58 | 12917 | Acidimicrobiia | Acidimicrobiales | Ilumatobacteraceae |  |  |
| Ilu_DEFall03L08_21 | 80.29 | 1.42 | 277 | 47996 | 1625841 | 1.63 | 59.95 | 7703 | Acidimicrobiia | Acidimicrobiales | Ilumatobacteraceae |  | Yes |
| Ilu_CPSum06L08_74 | 78.5 | 0 | 328 | 29192 | 2226312 | 2.23 | 64.98 | 9991 | Acidimicrobiia | Acidimicrobiales | Ilumatobacteraceae |  | Yes |
| Ilu_DESum019DL08_22 | 76.02 | 0 | 168 | 69449 | 1351996 | 1.35 | 50.12 | 15112 | Acidimicrobiia | Acidimicrobiales | Ilumatobacteraceae |  |  |
| Mic_DEBay_Spr20L08_14 | 94.88 | 0 | 34 | 165616 | 1220333 | 1.22 | 53.49 | 57454 | Actinomycetia | Actinomycetales | Microbacteriaceae | Yes |  |
| Mic_DESum22NL08_62 | 89.59 | 0 | 206 | 48840 | 1298501 | 1.30 | 56.89 | 9897 | Actinomycetia | Actinomycetales | Microbacteriaceae |  |  |
| Mic_CPSum06L08_5 | 87.56 | 1.52 | 114 | 71234 | 1148499 | 1.15 | 50.87 | 16344 | Actinomycetia | Actinomycetales | Microbacteriaceae |  |  |
| Mic_CPSpr15L08_24 | 83.1 | 0.58 | 85 | 73764 | 1144055 | 1.14 | 54.72 | 30070 | Actinomycetia | Actinomycetales | Microbacteriaceae |  | Yes |
| Mic_DEBay_Sum019D_LG_28 | 81.58 | 0.78 | 118 | 42562 | 1154121 | 1.15 | 47.65 | 14125 | Actinomycetia | Actinomycetales | Microbacteriaceae |  |  |
| Mic_CPSpr15G08_19 | 80.63 | 2.44 | 355 | 22505 | 1455364 | 1.46 | 58.66 | 5388 | Actinomycetia | Actinomycetales | Microbacteriaceae |  | Yes |
| Mic_DESum29DL08_23 | 79.65 | 1.66 | 236 | 48285 | 1068436 | 1.07 | 55.15 | 5675 | Actinomycetia | Actinomycetales | Microbacteriaceae | Yes |  |
| Mic_CPSum06L08_2 | 76.71 | 2.05 | 154 | 29733 | 1136986 | 1.14 | 48.09 | 10130 | Actinomycetia | Actinomycetales | Microbacteriaceae |  | Yes |
| Mil_DESum019DG08_62 | 91.38 | 0.86 | 140 | 80251 | 1741969 | 1.74 | 71.97 | 17765 | Thermoleophilia | Miltoncostaeales | Miltoncostaeaceae |  | Yes |
| Myc_DESum019DG08_34 | 94.89 | 0.6 | 355 | 72599 | 3615427 | 3.62 | 68 | 16927 | Actinomycetia | Mycobacteriales | Mycobacteriaceae |  |  |
| Nan_CPBay_Spr15L08_21 | 89.57 | 0.53 | 32 | 181136 | 1335996 | 1.34 | 47.81 | 73958 | Actinomycetia | Nanopelagicales | Nanopelagicaceae |  | Yes |
| Nan_CPSpr15L08_17 | 88.16 | 1.05 | 80 | 116484 | 1346422 | 1.35 | 45.08 | 35435 | Actinomycetia | Nanopelagicales | Nanopelagicaceae |  |  |
| Nan_CPSum15L08_12 | 86.1 | 0.75 | 78 | 117224 | 1230314 | 1.23 | 47.27 | 42311 | Actinomycetia | Nanopelagicales | Nanopelagicaceae | Yes | Yes |
| Nan_CPSum06L08_38 | 85.85 | 0.53 | 172 | 50968 | 1342913 | 1.34 | 51.67 | 10942 | Actinomycetia | Nanopelagicales | Nanopelagicaceae |  | Yes |
| Nan_CPSum15L08_1 | 84.4 | 0.68 | 97 | 59975 | 1155225 | 1.16 | 44.32 | 20826 | Actinomycetia | Nanopelagicales | Nanopelagicaceae | Yes |  |
| Nan_CPSum06L08_7 | 84.05 | 0.26 | 96 | 102794 | 1365676 | 1.37 | 50.09 | 22132 | Actinomycetia | Nanopelagicales | Nanopelagicaceae |  | Yes |
| Nan_DEFall03L08_2 | 83.41 | 3.07 | 277 | 35205 | 1374839 | 1.37 | 48.28 | 6303 | Actinomycetia | Nanopelagicales | Nanopelagicaceae |  | Yes |
| Nan_CPSpr15L08_54 | 82.78 | 1.05 | 68 | 92620 | 1199826 | 1.20 | 47.12 | 28439 | Actinomycetia | Nanopelagicales | Nanopelagicaceae |  | Yes |
| Nan_DEBay_Sum019D_LG_75 | 82.05 | 3.45 | 96 | 64259 | 1152602 | 1.15 | 48.19 | 18303 | Actinomycetia | Nanopelagicales | Nanopelagicaceae |  |  |
| Nan_DEFall15G08_39 | 80.45 | 0.53 | 80 | 129884 | 1140778 | 1.14 | 44.19 | 24502 | Actinomycetia | Nanopelagicales | Nanopelagicaceae |  | Yes |
| Nan_DESum019NG08_17 | 79.31 | 0 | 197 | 72879 | 1076409 | 1.08 | 43.81 | 6575 | Actinomycetia | Nanopelagicales | Nanopelagicaceae |  |  |
| Nan_DESum019DL08_13 | 78.73 | 1.4 | 91 | 65109 | 1192080 | 1.19 | 46.44 | 21376 | Actinomycetia | Nanopelagicales | Nanopelagicaceae |  | Yes |
| Nan_CPSum06G08_12 | 77.9 | 1.72 | 114 | 54343 | 777439 | 0.78 | 40.06 | 11257 | Actinomycetia | Nanopelagicales | Nanopelagicaceae |  |  |
| Nan_DESum019DL08_2 | 76.72 | 3.45 | 118 | 78917 | 1314387 | 1.31 | 46.74 | 18494 | Actinomycetia | Nanopelagicales | Nanopelagicaceae |  | Yes |
| Nan_CPSum06L08_80 | 76.43 | 0.08 | 165 | 57506 | 1252464 | 1.25 | 55.06 | 10818 | Actinomycetia | Nanopelagicales | Nanopelagicaceae |  |  |
| Nan_DESum019DL08_8 | 75.82 | 0 | 169 | 48492 | 1165930 | 1.17 | 43.88 | 10698 | Actinomycetia | Nanopelagicales | Nanopelagicaceae |  | Yes |
| Nan_DEFall03L08_12 | 75.43 | 1.72 | 118 | 64423 | 1117777 | 1.12 | 46.51 | 12368 | Actinomycetia | Nanopelagicales | Nanopelagicaceae |  | Yes |
| Nan_DESum019DL08_5 | 75 | 0 | 119 | 44751 | 917441 | 0.92 | 46.71 | 11992 | Actinomycetia | Nanopelagicales | Nanopelagicaceae |  | Yes |
| PAL_DESum019DG08_51 | 91.31 | 2.31 | 518 | 30102 | 2913112 | 2.91 | 72.74 | 7479 | Acidimicrobiia | IMCC26256 | PALSA-555 |  |  |
| S36_CPBay_Spr15L08_64 | 94.79 | 1.08 | 117 | 143459 | 2270042 | 2.27 | 60.04 | 29899 | Actinomycetia | Nanopelagicales | S36-B12 |  |  |
| S36_DEFall15L08_45 | 93.84 | 2.28 | 147 | 129537 | 2685904 | 2.69 | 66.07 | 36649 | Actinomycetia | Nanopelagicales | S36-B12 |  | Yes |
| S36_CPSpr30_3_6 | 93.57 | 0.54 | 103 | 129446 | 2047669 | 2.05 | 53.4 | 40013 | Actinomycetia | Nanopelagicales | S36-B12 |  |  |
| S36_DESpr20G08_17 | 92.67 | 0.54 | 94 | 172488 | 2350988 | 2.35 | 52.62 | 52339 | Actinomycetia | Nanopelagicales | S36-B12 |  | Yes |
| S36_DESum29DL08_29 | 92.54 | 0.77 | 233 | 48016 | 1982261 | 1.98 | 62.59 | 12175 | Actinomycetia | Nanopelagicales | S36-B12 |  | Yes |
| S36_CPSum06L08_10 | 89.93 | 1.13 | 145 | 111633 | 2447616 | 2.45 | 70 | 28936 | Actinomycetia | Nanopelagicales | S36-B12 |  | Yes |
| S36_DESum29NL08_47 | 89.82 | 0.96 | 405 | 32669 | 2147348 | 2.15 | 64.64 | 7015 | Actinomycetia | Nanopelagicales | S36-B12 |  |  |
| S36_CPSpr30_3_3 | 88.18 | 0.54 | 158 | 98045 | 1606619 | 1.61 | 53.49 | 16884 | Actinomycetia | Nanopelagicales | S36-B12 |  | Yes |
| S36_CPBay_Spr15L08_96 | 86.92 | 1.7 | 161 | 128454 | 2003769 | 2.00 | 52.31 | 21421 | Actinomycetia | Nanopelagicales | S36-B12 |  |  |
| S36_CPSum06L08_6 | 86.55 | 1.05 | 312 | 58169 | 2213411 | 2.21 | 67.71 | 10213 | Actinomycetia | Nanopelagicales | S36-B12 |  | Yes |
| S36_DESum29DL08_38 | 84.16 | 2.61 | 463 | 41225 | 2026154 | 2.03 | 62.25 | 5338 | Actinomycetia | Nanopelagicales | S36-B12 |  | Yes |
| S36_CPSpr30_6_4 | 80.1 | 4.74 | 484 | 36452 | 1953987 | 1.95 | 53.43 | 4802 | Actinomycetia | Nanopelagicales | S36-B12 |  | Yes |
| UB1_DEBay_Sum22DL08_24 | 94.87 | 2.56 | 118 | 144782 | 2318810 | 2.32 | 61.95 | 33145 | Acidimicrobiia | Acidimicrobiales | UBA11606 |  | Yes |
| UB5_DESum019DG08_56 | 79.36 | 0.54 | 309 | 32334 | 1611182 | 1.61 | 56.36 | 6589 | Actinomycetia | Nanopelagicales | UBA5976 |  | Yes |

**Table S2:** % of differentially expressed gene compared to all annotated gene in *Actinobacteriota* in various environmental conditions.

| Environmental Conditions | Ilu_CPSum15G08_28 | Nan_CPSum15L08_1 | Nan_CPSum15L08_12 | Mic_DEBay_Spr20L08_14 | Ilu_DEBay_Sum22DL08_50 | Ilu_DEBay_Sum29NL08_77 | Mic_DESum29DL08_23 | Median | Average |
| --- | --- | --- | --- | --- | --- | --- | --- | --- | --- |
| CPMidL08_SumvsSpr |  | 8.72 |  |  |  |  | 22.78 | 15.75 | 15.75 |
| CPSprL08_MidvsHi |  |  |  | 14.10 |  |  | 15.63 | 14.86 | 14.86 |
| CPSumG08_MidvsLow |  |  | 21.22 |  |  |  |  | 21.22 | 21.22 |
| CPSumL08_LowvsHi |  |  |  |  |  | 2.76 |  | 2.76 | 2.76 |
| CPSumL08_MidvsHi | 6.98 |  |  |  |  |  |  | 6.98 | 6.98 |
| CPSumL08_MidvsLow |  | 29.07 | 17.18 |  |  | 3.37 |  | 17.18 | 16.54 |
| CPSumLow_L08vsG08 |  |  | 11.81 |  |  |  |  | 11.81 | 11.81 |
| CPSumMid_L08vsG08 | 0.00 | 10.00 | 11.81 |  |  | 23.08 | 7.29 | 10.00 | 10.44 |
| DESumL08_MidvsHi | 0.23 |  |  |  | 0.05 | 1.29 |  | 0.23 | 0.52 |
| SprHiL08_DEvsCP |  |  |  | 39.02 |  |  |  | 39.02 | 39.02 |
| SumHiL08_DEvsCP | 9.37 |  |  |  |  | 1.80 | 3.42 | 3.42 | 4.86 |
| SumMidL08_DEvsCP | 8.74 |  |  |  | 3.52 | 8.62 | 42.31 | 8.68 | 15.80 |
| Median | 6.98 | 10.00 | 14.50 | 26.56 | 1.79 | 3.07 | 15.63 |  |  |
| Average | 5.06 | 15.93 | 15.51 | 26.56 | 1.79 | 6.82 | 18.29 |  |  |

## References

1. Traxler MF, Kolter R. 2015. Natural products in soil microbe interactions and evolution. Nat Prod Rep 32:956–970.

2. Zhang JW, Wang R, Liang X, Han P, Zheng YL, Li XF, Gao DZ, Liu M, Hou LJ, Dong HP. 2023. Novel Gene Clusters for Natural Product Synthesis Are Abundant in the Mangrove Swamp Microbiome. Appl Environ Microbiol 89.

3. Albarano L, Esposito R, Ruocco N, Costantini M. 2020. Genome mining as new challenge in natural products discovery. Mar Drugs 10.3390/md18040199.

4. Matsumoto A, Takahashi Y. 2017. Endophytic actinomycetes: Promising source of novel bioactive compounds. Journal of Antibiotics. Nature Publishing Group 10.1038/ja.2017.20.

5. Newton RJ, Jones SE, Eiler A, McMahon KD, Bertilsson S. 2011. A Guide to the Natural History of Freshwater Lake Bacteria. Microbiology and Molecular Biology Reviews 75:14–49.

6. Penn K, Jensen PR. 2012. Comparative genomics reveals evidence of marine adaptation in Salinispora species. BMC Genomics 13:1–12.

7. Ghai R, Mizuno CM, Picazo A, Camacho A, Rodriguez-Valera F. 2013. Metagenomics uncovers a new group of low GC and ultra-small marine Actinobacteria. Sci Rep 3:1–8.

8. Udwary DW, Zeigler L, Asolkar RN, Singan V, Lapidus A, Fenical W, Jensen PR, Moore BS. 2007. Genome sequencing reveals complex secondary metabolome in the marine actinomycete Salinispora tropica. Proc Natl Acad Sci U S A 104:10376–10381.

9. Mehrshad M, Amoozegar MA, Ghai R, Shahzadeh Fazeli SA, Rodriguez-Valera F. 2016. Genome reconstruction from metagenomic data sets reveals novel microbes in the brackish waters of the Caspian Sea. Appl Environ Microbiol 82:1599–1612.

10. Prabhu A, Tule S, Chuvochina M, Bodén M, Mcilroy SJ, Zaugg J, Rinke C. Machine learning and metagenomics identifies uncharacterized taxa inferred to drive biogeochemical cycles in a subtropical hypereutrophic estuary. ISME Communications 2024:67.

11. Barka EA, Vatsa P, Sanchez L, Gaveau-Vaillant N, Jacquard C, Klenk H-P, Clément C, Ouhdouch Y, van Wezel GP. 2016. Taxonomy, Physiology, and Natural Products of Actinobacteria. Microbiology and Molecular Biology Reviews 80:1–43.

12. Ghai R, Mizuno CM, Picazo A, Camacho A, Rodriguez-Valera F. 2014. Key roles for freshwater Actinobacteria revealed by deep metagenomic sequencing. Mol Ecol 23:6073–6090.

13. Ghai R, Mcmahon KD, Rodriguez-Valera F. 2012. Breaking a paradigm: Cosmopolitan and abundant freshwater actinobacteria are low GC. Environ Microbiol Rep 4:29–35.

14. Holmfeldt K, Dziallas C, Titelman J, Pohlmann K, Grossart HP, Riemann L. 2009. Diversity and abundance of freshwater Actinobacteria along environmental gradients in the brackish northern Baltic Sea. Environ Microbiol 11:2042–2054.

15. Bérdy J. 2005. Bioactive microbial metabolites: A personal view. Journal of Antibiotics. Japan Antibiotics Research Association 10.1038/ja.2005.1.

16. Undabarrena A, Beltrametti F, Claverías FP, González M, Moore ERB, Seeger M, Cámara B. 2016. Exploring the diversity and antimicrobial potential of marine actinobacteria from the comau fjord in Northern Patagonia, Chile. Front Microbiol 7:1135.

17. Montagna PA, Palmer TA, Beseres Pollack J. 2013. Conceptual Model of Estuary Ecosystems, p. 5–21. *In* Hydrological Changes and Estuarine Dynamics. Springer New York, New York, NY.

18. Telesh I V., Khlebovich V V. 2010. Principal processes within the estuarine salinity gradient: A review. Mar Pollut Bull 61:149–155.

19. Kan J, Crump BC, Wang K, Chen F. 2006. Bacterioplankton community in Chesapeake Bay: Predictable or random assemblages. Limnol Oceanogr 51:2157–2169.

20. Cram JA, Hollins A, McCarty AJ, Martinez G, Cui M, Gomes ML, Fuchsman CA. 2024. Microbial diversity and abundance vary along salinity, oxygen, and particle size gradients in the Chesapeake Bay. Environ Microbiol 26:e16557.

21. Selje N, Simon M. 2003. Composition and dynamics of particle-associated and free-living bacterial communities in the Weser estuary, Germany. Aquatic Microbial Ecology 30:221–237.

22. Leu AO, Eppley JM, Burger A, DeLong EF, Maria Gloria Dominguez Bello E, Jan-Hedrik Hehemanm by, Tully B. 2025. Diverse genomic traits differentiate sinking-particle-associated versus free-living microbes throughout the oligotrophic open ocean water column. journals.asm.org AO Leu, JM Eppley, A Burger, EF DeLong MBio, 2022•journals.asm.org 13.

23. Kirchman DL, Dittel AI, Malmstrom RR, Cottrell MT. 2005. Biogeography of major bacterial groups in the Delaware Estuary. Limnol Oceanogr 50:1697–1706.

24. Sarin MM, Church TM. 1994. Behaviour of uranium during mixing in the delaware and chesapeake estuaries. Estuar Coast Shelf Sci 39:619–631.

25. Toschik PC, Rattner BA, McGowan PC, Christman MC, Carter DB, Hale RC, Matson CW, Ottinger MA. 2005. EFFECTS OF CONTAMINANT EXPOSURE ON REPRODUCTIVE SUCCESS OF OSPREYS (PANDION HALIAETUS) NESTING IN DELAWARE RIVER AND BAY, USA. Environ Toxicol Chem 24:617.

26. Kanehisa M, Sato Y, Morishima K. 2016. BlastKOALA and GhostKOALA: KEGG Tools for Functional Characterization of Genome and Metagenome Sequences. J Mol Biol 428:726–731.

27. Blin K, Shaw S, Kloosterman AM, Charlop-Powers Z, Van Wezel GP, Medema MH, Weber T. 2021. antiSMASH 6.0: improving cluster detection and comparison capabilities. Nucleic Acids Res 49:W29–W35.

28. Finn RD, Attwood TK, Babbitt PC, Bateman A, Bork P, Bridge AJ, Chang HY, Dosztanyi Z, El-Gebali S, Fraser M, Gough J, Haft D, Holliday GL, Huang H, Huang X, Letunic I, Lopez R, Lu S, Marchler-Bauer A, Mi H, Mistry J, Natale DA, Necci M, Nuka G, Orengo CA, Park Y, Pesseat S, Piovesan D, Potter SC, Rawlings ND, Redaschi N, Richardson L, Rivoire C, Sangrador-Vegas A, Sigrist C, Sillitoe I, Smithers B, Squizzato S, Sutton G, Thanki N, Thomas PD, Tosatto SCE, Wu CH, Xenarios I, Yeh LS, Young SY, Mitchell AL. 2017. InterPro in 2017—beyond protein family and domain annotations. Nucleic Acids Res 45:D190–D199.

29. Jones P, Binns D, Chang HY, Fraser M, Li W, McAnulla C, McWilliam H, Maslen J, Mitchell A, Nuka G, Pesseat S, Quinn AF, Sangrador-Vegas A, Scheremetjew M, Yong SY, Lopez R, Hunter S. 2014. InterProScan 5: genome-scale protein function classification. Bioinformatics 30:1236–1240.

30. Joseph TA, Chlenski P, Litman A, Korem T, Pe’er I. 2022. Accurate and robust inference of microbial growth dynamics from metagenomic sequencing reveals personalized growth rates. Genome Res 32:558–568.

31. Campbell BJ, Kirchman DL. 2013. Bacterial diversity, community structure and potential growth rates along an estuarine salinity gradient. The ISME Journal 2013 7:1 7:210–220.

32. Wu Z, Li M, Qu L, Zhang C, Xie W. 2024. Metagenomic insights into microbial adaptation to the salinity gradient of a typical short residence-time estuary. Microbiome 12:1–19.

33. Zakharova Y, Bashenkhaeva M, Galachyants Y, Petrova D, Tomberg I, Marchenkov A, Kopyrina L, Likhoshway Y. 2022. Variability of Microbial Communities in Two Long-Term Ice-Covered Freshwater Lakes in the Subarctic Region of Yakutia, Russia. Microb Ecol 84:958–973.

34. Silva-Solar S, Viver T, Wang Y, Orellana LH, Knittel K, Amann R. 2024. Acidimicrobiia, the actinomycetota of coastal marine sediments: Abundance, taxonomy and genomic potential. Syst Appl Microbiol 47:126555.

35. Campbell BJ, Yu L, Heidelberg JF, Kirchman DL. 2011. Activity of abundant and rare bacteria in a coastal ocean. Proc Natl Acad Sci U S A 108:12776–12781.

36. Rudolf JD, Alsup TA, Xu B, Li Z. 2021. Bacterial terpenome. Nat Prod Rep 38:905–980.

37. Avalos M, Garbeva P, Vader L, Wezel GP Van, Dickschat JS, Ulanova D. 2022. Biosynthesis, evolution and ecology of microbial terpenoids. Nat Prod Rep 39:249–272.

38. Rego A, Fernandez-Guerra A, Duarte P, Assmy P, Leão PN, Magalhães C. 2021. Secondary metabolite biosynthetic diversity in Arctic Ocean metagenomes. Microb Genom 7:731.

39. Zhang Z, Zhang L, Zhang L, Chu H, Zhou J, Ju F. 2024. Diversity and distribution of biosynthetic gene clusters in agricultural soil microbiomes. mSystems 9.

40. Waschulin V, Borsetto C, James R, Newsham KK, Donadio S, Corre C, Wellington E. ARTICLE Biosynthetic potential of uncultured Antarctic soil bacteria revealed through long-read metagenomic sequencing 10.1038/s41396-021-01052-3.

41. Sánchez-Navarro R, Nuhamunada M, Mohite OS, Wasmund K, Albertsen M, Gram L, Nielsen PH, Weber T, Singleton CM. 2022. Long-Read Metagenome-Assembled Genomes Improve Identification of Novel Complete Biosynthetic Gene Clusters in a Complex Microbial Activated Sludge Ecosystem. mSystems 7.

42. Tarasova E V., Luchnikova NA, Grishko V V., Ivshina IB. 2023. Actinomycetes as Producers of Biologically Active Terpenoids: Current Trends and Patents. Pharmaceuticals 16:872.

43. Rudolf JD, Alsup TA, Xu B, Li Z. 2021. Bacterial terpenome. Nat Prod Rep 38:905–980.

44. Bruna P, Núñez-Montero K, Contreras MJ, Leal K, García M, Abanto M, Barrientos L. 2024. Biosynthetic gene clusters with biotechnological applications in novel Antarctic isolates from Actinomycetota. Appl Microbiol Biotechnol 108:1–11.

45. Chen Y, Yang Y, Ji X, Zhao R, Li G, Gu Y, Shi A, Jiang W, Zhang Q. 2020. The SCIFF-Derived Ranthipeptides Participate in Quorum Sensing in Solventogenic Clostridia. Biotechnol J 15:2000136.

46. Zhang Z, Zhang L, Zhang L, Chu H, Zhou J, Ju F. 2024. Diversity and distribution of biosynthetic gene clusters in agricultural soil microbiomes. mSystems 9.

47. Fisher TR, Harding LW, Stanley DW, Ward LG. 1988. Phytoplankton, nutrients, and turbidity in the Chesapeake, Delaware, and Hudson estuaries. Estuar Coast Shelf Sci 27:61–93.

48. Umanskaya M V., Gorbunov MY. 2024. Phylogenetic Structure of Bacterioplankton in Water Bodies of the Kuibyshev Reservoir Basin during Cyanobacterial Bloom. Microbiology (Russian Federation) 93:876–890.

49. Chiriac M-C, Haber M, Salcher MM. 2022. Adaptive genetic traits in pelagic freshwater microbes. Wiley Online Library MC Chiriac, M Haber, MM SalcherEnvironmental Microbiology, 2023•Wiley Online Library 25:606–641.

50. Rohwer RR, Kirkpatrick M, Garcia SL, Kellom M, McMahon KD, Baker BJ. 2025. Two decades of bacterial ecology and evolution in a freshwater lake. Nat Microbiol 10:246–257.

51. Wu Z, Li M, Qu L, Zhang C, Xie W. 2024. Metagenomic insights into microbial adaptation to the salinity gradient of a typical short residence-time estuary. Microbiome 12:1–19.

52. Seidel L, Broman E, Ståhle M, Bergström K, Forsman A, Hylander S, Ketzer M, Dopson M. 2024. Climate change induces shifts in coastal Baltic Sea surface water microorganism stress and photosynthesis gene expression. Front Microbiol 15:1393538.

53. Ghai R, Mizuno C, Picazo A, … AC-M, 2014 undefined. 2014. Key roles for freshwater A ctinobacteria revealed by deep metagenomic sequencing. Wiley Online Library R Ghai, CM Mizuno, A Picazo, A Camacho, F Rodriguez-Valera Molecular ecology, 2014•Wiley Online Library 23:6073–6090.

54. Neuenschwander SM, Ghai R, Pernthaler J, Salcher MM. 2018. Microdiversification in genome-streamlined ubiquitous freshwater Actinobacteria. academic.oup.comSM Neuenschwander, R Ghai, J Pernthaler, MM SalcherThe ISME journal, 2018•academic.oup.com 10.1038/ismej.2017.156.

55. Galasso C, Corinaldesi C, Sansone C. 2017. Carotenoids from Marine Organisms: Biological Functions and Industrial Applications. Antioxidants 2017, Vol 6, Page 96 6:96.

56. Maresca JA, Keffer JL, Hempel PP, Polson SW, Shevchenko O, Bhavsar J, Powell D, Miller KJ, Singh A, Hahn MW. 2019. Light modulates the physiology of nonphototrophic Actinobacteria. J Bacteriol 209:740–758.

57. Ottesen EA, Young CR, Eppley JM, Ryan JP, Chavez FP, Scholin CA, DeLong EF. 2013. Pattern and synchrony of gene expression among sympatric marine microbial populations. Proceedings of the National Academy of Sciences 110:E488–E497.

58. Ahmed MA, Campbell BJ. 2025. Genome-resolved adaptation strategies of Rhodobacterales to changing conditions in the Chesapeake and Delaware Bays. Appl Environ Microbiol 91.

59. Stevens H, Brinkhoff T, Rink B, Vollmers J, Simon M. 2007. Diversity and abundance of Gram positive bacteria in a tidal flat ecosystem. Environ Microbiol 9:1810–1822.

60. Ayayee PA, Custer GF, Tronstad LM, van Diepen LTA. 2024. Unveiling salinity-driven shifts in microbial community composition across compartments of naturally saline inland streams. Hydrobiologia 851:2627–2639.

61. Pitt A, Schmidt J, Koll U, Hahn MW. 2021. Aquiluna borgnonia gen. Nov., sp. nov., a member of a microbacteriaceae lineage of freshwater bacteria with small genome sizes. Int J Syst Evol Microbiol 71:004825.

62. Han SK, Nedashkovskaya OI, Mikhailov V V., Kim SB, Bae KS. 2003. Salinibacterium amurskyense gen. nov., sp. nov., a novel genus of the family Microbacteriaceae from the marine environment. Int J Syst Evol Microbiol 53:2061–2066.

63. Stackebrandt E, Brambilla E, Richert K. 2007. Gene sequence phylogenies of the family Microbacteriaceae. Curr Microbiol 55:42–46.

64. Hu WJ, Deng LX, Huang YY, Wang XC, Qing JL, Zhu HJ, Zhou X, Zhou XY, Chu JM, Pan X. 2025. Genome mining and metabolite profiling illuminate the taxonomy status and the cytotoxic activity of a mangrove-derived Microbacterium alkaliflavum sp. nov. BMC Microbiol 25:1–13.

65. Gupta S, Han SR, Kim B, Lee CM, Oh TJ. 2022. Comparative analysis of genome-based CAZyme cassette in Antarctic Microbacterium sp. PAMC28756 with 31 other Microbacterium species. Genes Genomics 44:733–746.

66. Jeffries TC, Schmitz Fontes ML, Harrison DP, Van-Dongen-Vogels V, Eyre BD, Ralph PJ, Seymour JR. 2016. Bacterioplankton dynamics within a large anthropogenically impacted urban estuary. Front Microbiol 6:163345.

67. Smith MW, Herfort L, Tyrol K, Suciu D, Campbell V, Crump BC, Peterson TD, Zuber P, Baptista AM, Simon HM. 2010. Seasonal Changes in Bacterial and Archaeal Gene Expression Patterns across Salinity Gradients in the Columbia River Coastal Margin. PLoS One 5:e13312.

68. Chase AB, Weihe C, Martiny JBH. 2021. Adaptive differentiation and rapid evolution of a soil bacterium along a climate gradient. Proceedings of the National Academy of Sciences 118:e2101254118.

69. Chase AB, Karaoz U, Brodie EL, Gomez-Lunar Z, Martiny AC, Martiny JBH. 2017. Microdiversity of an abundant terrestrial bacterium encompasses extensive variation in ecologically relevant traits. mBio 8.

70. Bech PK, Jarmusch SA, Rasmussen JA, Limborg MT, Gram L, Henriksen NNSE. 2024. Succession of microbial community composition and secondary metabolism during marine biofilm development. ISME Communications 4:6.

71. Marchese P, Bracegirdle J, Young R, Ferrari E, Garzoli L, Murphy JM, Tuohy M, Allcock AL, Baker BJ. 2025. Bacterial Diversity in Deep-Sea Sediments of the North Atlantic Ocean and Their Biosynthesis of Secondary Metabolites. Environ Microbiol Rep 17:e70092.

72. Corretto E, Antonielli L, Sessitsch A, Höfer C, Puschenreiter M, Widhalm S, Swarnalakshmi K, Brader G. 2020. Comparative Genomics of Microbacterium Species to Reveal Diversity, Potential for Secondary Metabolites and Heavy Metal Resistance. Front Microbiol 11:531937.

73. Cordovez V, Schop S, Hordijk K, de Boulois HD, Coppens F, Hanssen I, Raaijmakers JM, Carrióna VJ. 2018. Priming of plant growth promotion by volatiles of rootassociated Microbacterium spp. Appl Environ Microbiol 84:1865–1883.

74. Qiao LK, He LY, Gao FZ, Huang Z, Bai H, Wang YC, Shi YJ, Liu YS, Zhao JL, Ying GG. 2025. Deciphering key traits and dissemination of antibiotic resistance genes and degradation genes in pharmaceutical wastewater receiving environments. Water Res 275:123241.

75. Goodarzi Z, Asad S, Mehrshad M. 2022. Genome-resolved insight into the reservoir of antibiotic resistance genes in aquatic microbial community. Scientific Reports 2022 12:1 12:1–11.

76. Freund L, Hung C, Topacio TM, Diamond C, Fresquez A, Lyons TW, Aronson EL. 2025. Diversity of sulfur cycling halophiles within the Salton Sea, California’s largest lake. BMC Microbiol 25:1–24.

77. Ahmed MA, Lim SJ, Campbell BJ. 2021. Metagenomes, Metatranscriptomes, and Metagenome-Assembled Genomes from Chesapeake and Delaware Bay (USA) Water Samples. Microbiol Resour Announc 10.

78. Maresca JA, Miller KJ, Keffer JL, Sabanayagam CR, Campbell BJ. 2018. Distribution and diversity of rhodopsinproducing microbes in the Chesapeake Bay. Appl Environ Microbiol 84.

79. Gurevich A, Saveliev V, Vyahhi N, Tesler G. 2013. QUAST: Quality assessment tool for genome assemblies. Bioinformatics 29:1072–1075.

80. Parks DH, Imelfort M, Skennerton CT, Hugenholtz P, Tyson GW. 2015. CheckM: Assessing the quality of microbial genomes recovered from isolates, single cells, and metagenomes. Genome Res 25:1043–1055.

81. Olm MR, Brown CT, Brooks B, Banfield JF. 2017. DRep: A tool for fast and accurate genomic comparisons that enables improved genome recovery from metagenomes through de-replication. ISME Journal 11:2864–2868.

82. Parks DH, Chuvochina M, Waite DW, Rinke C, Skarshewski A, Chaumeil PA, Hugenholtz P. 2018. A standardized bacterial taxonomy based on genome phylogeny substantially revises the tree of life. Nat Biotechnol 36:996.

83. Letunic I, Bork P. 2021. Interactive Tree Of Life (iTOL) v5: an online tool for phylogenetic tree display and annotation. Nucleic Acids Res 49:W293–W296.

84. Langmead B, Salzberg SL. 2012. Fast gapped-read alignment with Bowtie 2. Nat Methods 9:357–359.

85. Li H, Handsaker B, Wysoker A, Fennell T, Ruan J, Homer N, Marth G, Abecasis G, Durbin R, Project G, Subgroup DP. 2009. The Sequence Alignment/Map format and SAMtools 25:2078–2079.

86. Menzel P, Ng KL, Krogh A. 2016. Fast and sensitive taxonomic classification for metagenomics with Kaiju. Nature Communications 2016 7:1 7:1–9.

87. Wickham MH. 2014. Package “ggplot2” Type Package Title An implementation of the Grammar of Graphics.

88. Team RC. 2022. R: A Language and Environment for Statistical Computing.

89. Gu Z. 2022. Complex heatmap visualization. iMeta 1:e43.

90. Wickham H. 2016. ggplot2 10.1007/978-3-319-24277-4.

91. Korem T, Zeevi D, Suez J, Weinberger A, Avnit-Sagi T, Pompan-Lotan M, Matot E, Jona G, Harmelin A, Cohen N, Sirota-Madi A, Thaiss CA, Pevsner-Fischer M, Sorek R, Xavier RJ, Elinav E, Segal E. 2015. Growth dynamics of gut microbiota in health and disease inferred from single metagenomic samples. Science (1979) 349:1101–1106.

92. Galili T, O’Callaghan A, Sidi J, Sievert C. 2018. Heatmaply: An R package for creating interactive cluster heatmaps for online publishing. Bioinformatics 34:1600–1602.

93. Seemann T. 2014. Genome analysis Prokka: rapid prokaryotic genome annotation 30:2068–2069.

94. Medema MH, Blin K, Cimermancic P, De Jager V, Zakrzewski P, Fischbach MA, Weber T, Takano E, Breitling R. 2011. antiSMASH: rapid identification, annotation and analysis of secondary metabolite biosynthesis gene clusters in bacterial and fungal genome sequences. Nucleic Acids Res 39:W339–W346.

95. Mungan MD, Alanjary M, Blin K, Weber T, Medema MH, Ziemert N. 2020. ARTS 2.0: feature updates and expansion of the Antibiotic Resistant Target Seeker for comparative genome mining. Nucleic Acids Res 48:W546–W552.

96. Eren AM, Esen OC, Quince C, Vineis JH, Morrison HG, Sogin ML, Delmont TO. 2015. Anvi’o: An advanced analysis and visualization platformfor ‘omics data. PeerJ 2015:e1319.

97. Liao Y, Smyth GK, Shi W. 2014. featureCounts: an efficient general purpose program for assigning sequence reads to genomic features. Bioinformatics 30:923–930.

98. Kanehisa M, Goto S. 2000. KEGG: Kyoto Encyclopedia of Genes and Genomes. Nucleic Acids Res 28:27–30.

99. Love MI, Huber W, Anders S. 2014. Moderated estimation of fold change and dispersion for RNA-seq data with DESeq2. Genome Biol 15.

100. Chaumeil PA, Mussig AJ, Hugenholtz P, Parks DH. 2020. GTDB-Tk: A toolkit to classify genomes with the genome taxonomy database. Bioinformatics 36:1925–1927.

